# Independent Lateralization of Language, Attention, and Numerical Cognition Across Task and Rest

**DOI:** 10.1101/2025.11.23.690045

**Authors:** Loïc Labache, Isabelle Hesling, Laure Zago

## Abstract

Hemispheric functional complementarity is a core organizational principle of the human brain, yet the extent to which lateralization in one domain constrains that of others remains unclear. Two main accounts have been proposed: the causal hypothesis, in which dominance for one function drives complementary dominance in another, and the statistical hypothesis, in which each function lateralizes independently. Using multimodal fMRI in 287 participants from the BIL&GIN cohort, we examined whether language lateralization phenotypes, defined as typical (left-dominant) or atypical (right-dominant), predict hemispheric asymmetries in visuospatial attention and numerical cognition. Task-based activation was measured during line bisection, mental calculation, and numerical interval comparison, and analyzed within domain-specific, functionally defined network atlases. Resting-state functional connectivity metrics were also assessed in the same networks. Across both attention and numerical domains, typical individuals for language showed stronger asymmetries, whereas atypical individuals exhibited weaker, more bilateral patterns. Critically, atypical participants did not show mirror-reversed asymmetries, and language phenotype did not influence intrinsic connectivity metrics in non-language networks. These findings challenge the notion that atypical lateralization represents an inversion of the canonical template and argue against a universal reciprocal link between language dominance and other cognitive domains. Instead, our results support a domain-specific model in which lateralization profiles are shaped by distinct developmental and functional constraints, highlighting the need for multimodal, multi-domain approaches to brain asymmetry.

---

**H**emispheric specialization enhances neural efficiency by reducing redundancy and enabling parallel, domain-specific processing (1–3). Classic examples include left-hemispheric dominance for language and praxis (4–8), and right-hemispheric dominance for visuospatial attention (9–13). While such patterns are well established within individual domains, it remains unclear whether the lateralization of one function shapes that of others. Two competing accounts have been proposed. The causal hypothesis predicts that dominance for one function drives complementary dominance for another; for example, left-dominant language constraining attention to the right hemisphere (14). The statistical hypothesis posits that each function lateralizes independently, with any cross-domain associations emerging probabilistically (15). Discriminating between these accounts is critical for understanding the developmental and neurobiological bases of brain organization. Under this view, atypical (rightward) language dominance should be accompanied by mirror-reversed asymmetries in visuospatial attention and numerical cognition, reflecting a global redistribution of functions across hemispheres. In contrast, the statistical hypothesis predicts that each domain lateralizes independently, such that cross-domain associations arise only from probabilistic correlations at the population level. This account predicts that atypical language dominance should not systematically influence the direction, magnitude, or intrinsic connectivity of asymmetries in non-language networks. Our multimodal design allows us to directly discriminate between these predictions by assessing whether language phenotype explains interindividual variation in both task-evoked and intrinsic hemispheric organization across attention and numerical domains.

While these canonical patterns are well established, even within a single domain such as language, lateralization can vary. Left-lateralized fronto-temporal circuits support syntactic and grammatical processing, whereas right-lateralized regions contribute to prosody and contextual interpretation (16, 17). In visuospatial attention, the temporo-frontal network encompasses both rightward temporal-frontal regions and leftward superior temporal cortex and subcortical nuclei, suggesting a broader functional scope than previously recognized (18). A similar hemispheric division of labor is seen in numerical cognition: symbolic arithmetic operations and rule-based manipulations exhibit a left-hemispheric bias, whereas approximate number estimation and non-symbolic magnitude comparison are more robustly associated with right-hemispheric networks (19–22). These patterns of co-localization and segregation suggest that functional complementarity serves as a principle of cortical economy, whereby language and arithmetic converge within the left hemisphere, while spatial attention and magnitude estimation are preferentially instantiated in the right hemisphere.

Despite this well-established inter-domain asymmetry, the extent to which lateralization in one domain constrains, co-develops with, or is independent from other domains remains unresolved (23). Language, particularly speech production, has traditionally served as a reference point for classifying atypical lateralization across domains, given its early developmental onset and strong left-hemispheric bias (24). This assumption predicts that atypical language dominance should be accompanied by mirrored asymmetries in other domains, especially in individuals with non-leftward profiles. However, empirical evidence for such mirrored relationships is limited, particularly for numerical cognition, where symbolic and analog systems rely on distinct lateralized networks and may interact differently with language-related circuits (21, 25).

The developmental trajectory of hemispheric specialization suggests a gradual, multifactorial process shaped by genetic, epigenetic, and environmental influences (26). Evolutionary perspectives posit that lateralization evolved to enhance efficiency by distributing high-demand cognitive operations across hemispheres (27). Evidence for such segregation exists at multiple levels, macrostructural (e.g., corpus callosum connectivity (28)), microstructural and molecular (29), and large-scale network organization (16–18, 30, 31). Clinically, atypical lateralization patterns have been linked to neurodevelopmental and neuropsychiatric conditions including dyslexia, schizophrenia, and post-stroke recovery (32).

Previous neuroimaging studies have largely examined language, attention, and numerical networks in isolation. Yet emerging evidence suggests that hemispheric dominance may reflect broader patterns of interhemispheric coordination, with variability in one domain potentially related to variability in others. For instance, Gerrits and colleagues assessed lateralization for five functions in a sample of 63 individuals, enriched for right-hemisphere language dominance, using cytoarchitectonic region-of-interest analyses (33). They reported that right-hemisphere language dominance was frequently accompanied by reversed or nearly reversed lateralization of other functions, consistent with a strong cross-domain coupling. These findings suggest that, at least in some populations, hemispheric organization can be globally mirrored rather than independently determined, thereby lending support to a causal account of hemispheric complementarity. In addition, co-lateralization analyses have shown that language and symbolic number processing can covary in asymmetry profiles, particularly in frontoparietal regions (21). However, another study has reported weaker or inconsistent cross-domain associations, indicating that both interdependence and independence may coexist (34). Beyond task-evoked activations, resting-state functional connectivity offers insight into intrinsic organization. Typical leftward language dominance has been associated with leftward asymmetries in degree centrality, reduced interhemispheric homotopy, and lower global integration, whereas individuals with atypical dominance exhibit more symmetrical connectivity (35, 36). Whether such intrinsic markers generalize to other lateralized domains is unknown. Addressing this question is essential for determining whether hemispheric specialization arises from shared neurobiological constraints or function-specific developmental pathways.

Here, we test these alternatives by combining task-based and resting-state fMRI in a large, well-characterized sample of 287 participants from the BIL&GIN cohort (37). We examine to what extent language lateralization phenoty pepredictshemisphericasymmetries in visuospatial attention and numerical cognition networks, and whether such associations extend to intrinsic connectivity metrics. By integrating multi-domain, multimodal measures, we directly evaluate the causal and statistical hypotheses of hemispheric complementarity.

## Results

### Lateralized Brain Networks Supporting Visuospatial Attention (ALANs) and Numerical Cognition (LUCA) in right-handed left-language lateralized individuals

To evaluate how language lateralization phenotype shapes task-related asymmetry in visuospatial attention and numerical cognition networks, we first selected the lateralized networks associated with visuospatial attention, arithmetic, and number comparison in a sample of right-handed individuals typically left-lateralized for language (see Table 1 for sample characteristics).

**Table 1.** Demographic characteristics of the participants subsamples included in the atlas construction, task-based fMRI, and resting-state fMRI analyses for each cognitive domain. ALANs refer to the Atlas for Lateralized visuospatial Attentional Networks identified during the line bisection judgment task (Fig. 1a, (18)), and LUCA refers to the Lateralized Underpinnings of Comparison and Arithmetic networks atlas identified during a calculation and a numerical interval comparison tasks (Fig. 1b, Table S1). The table reports, for each subsample, the number of participants, the number of lefthanders (based on self-reported manual preference), the number of women, the number of participants with atypical language lateralization (*i*.*e*., right-hemisphere dominant), and the number of left-hand responses recorded during the line bisection task. All participants were drawn from the BIL&GIN database (37) and previously characterized for language lateralization using multimodal fMRI (35).

| Cognitive Function | Modality | Participant # | Manual Preference (left-handers #) | Gender (♀ #) | Language Lateralization Phenotype (atypical #) | Responding Hand (left-hand responses #) |
| --- | --- | --- | --- | --- | --- | --- |
| ALANs | atlas | 130 | 0 | 64 | 0 | 0 |
|  | task-fMRI | 284 | 149 | 139 | 30 | 18 |
|  | rs-fMRI | 284 | 149 | 139 | 30 | – |
| LUCA | atlas | 120 | 0 | 58 | 0 | 0 |
|  | task-fMRI | 250 | 126 | 122 | 25 | 13 |
|  | rs-fMRI | 287 | 150 | 140 | 30 | – |

For visuospatial attention, we relied on the Atlas for the Lateralized visuospatial Attention Networks (ALANs (18)), identified during the line-bisection judgment task. ALANs comprises five lateralized large-scale networks; parieto-frontal, temporo-frontal, posterior-medial, somato-motor, and visual, each defined by a weighted BOLD asymmetry procedure that quantifies task-evoked lateralization by comparing voxel-weighted activation in atlas-defined dominant-hemisphere regions to their homotopic counterparts. As illustrated in Fig. 1a, three networks are fully right-lateralized, one fully left-lateralized, and one, the temporo-frontal, predominantly right-lateralized with a few left-lateralized regions.

**Figure 1.**
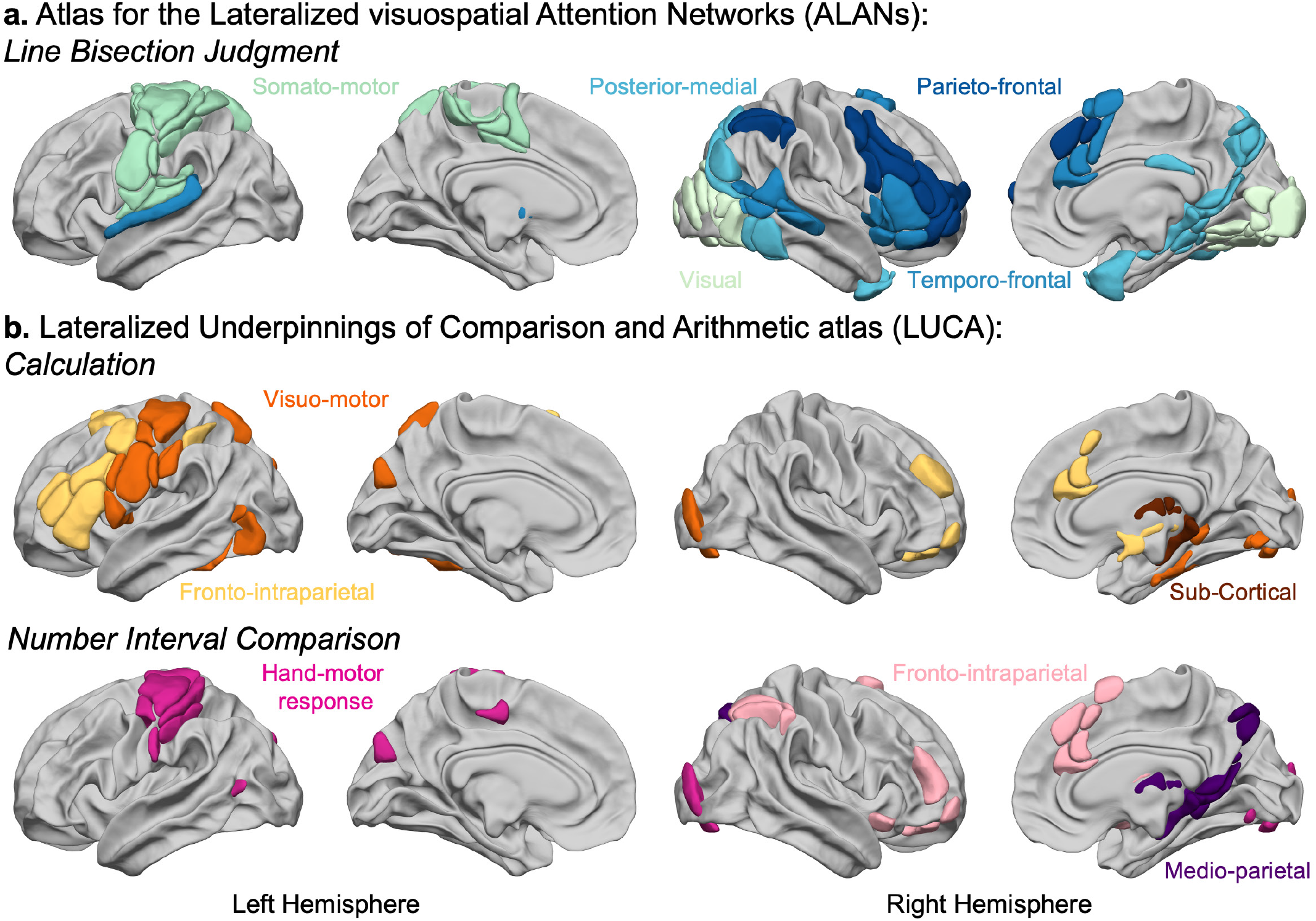
Lateralized Brain Networks Supporting Visuospatial Attention (ALANs), Calculation, and Numerical Interval comparison (LUCA : Lateralized Underpinnings of Comparison and Arithmetic atlas) in typical language-lateralized participants. **a**, Atlas for the lateralized visuospatial attention networks (ALANs). The atlas for the lateralized visuospatial attention networks comprises five networks that provide the anatomo-functional support for visual-spatial attention processing (18). This atlas includes two primary core attentional networks: the parieto-frontal and temporo-frontal networks, as well as three support networks: the posteriormedial network, the somato-motor, and the visual networks, which all support sensory-motor functions during visuospatial attention tasks. **b**, Top Row: Atlas for the lateralized underpinnings for the calculation task (see Table S1 for a full description of each region). This atlas includes three arithmetic networks: the fronto-intraparietal, the visuo-motor and subcortical networks. Bottom Row: Lateralized Underpinnings of numerical interval comparison task (see Table S1 for a full description of each region). This atlas includes three comparison networks: right fronto-intraparietal, right medioparietal and left hand-motor networks.

Using an analogous methodological framework (16–18), we identified the Lateralized Underpinnings of Comparison and Arithmetic (LUCA) atlas to characterize numerical cognition networks. LUCA was derived from regions showing significant leftward or rightward activation and asymmetry during calculation or numerical comparison in a large reference group of typically left-lateralized participants for language, followed by resting-state connectivity analyses to define task-specific network organization. As shown in Fig. 1b, calculation engaged three networks (fronto-intraparietal, visuo-motor, and subcortical), whereas comparison also recruited three networks (fronto-intraparietal, medio-parietal, and hand-motor response), each exhibiting characteristic leftward or rightward lateralization profiles.

### Impact of Language Lateralization Phenotypes on Visuospatial Attention Networks Asymmetries

To assess the effect of language lateralization phenotype on the task-related asymmetry of visuospatial attention networks, we conducted separate linear regression analyses for each of the five ALANs networks (Fig. 1).These analyses were performed on a sample of 284 participants from the BIL&GIN database (38), all having performed a line bisection judgment task (39), and whose language lateralization phenotype (typical left-lateralised, atypical right-lateralised) had been previously identified using multimodal language task-based fMRI (35). Weighted BOLD asymmetry scores during the task were the dependent variable, language lateralization phenotype the main predictor of interest, while controlling for manual preference, age, years of education, total intracranial volume, gender, response hand, and the interaction between language lateralization phenotype and manual preference (Fig. S1).

Type III analysis of covariance revealed a significant main effect of language lateralization phenotype on the asymmetry scores of three out of the five networks: the parieto-frontal (*p*_model FDR_=4.10^-4^, *p*^Phenotype FDR^=8.10^-3^, *η*^2^ _p_ =0.031), temporo-frontal (*p* _model FDR_ =6.10^-9^, *p*_Phenotype FDR_ =5.10^-4^, *η*^2^ _p_ =0.054), and visualnetworks (*p*_model FD R_=1.10^-3^, *p* _PhenotypeFDR_=2.10^-2^, *η*^2^ _p_=0.021) (Table 2, Fig. 2). No significant effects were observed on the asymmetry scores for the posterior-medial (*p*_model FDR_=5.10^-2^, *p*_Phenotype FDR_=2.10^-2^, *η*^2^_p_=0.023) or the somato-motor networks (*p*_modelFDR_=4.10^-38^, *p* _PhenotypeFDR_=4.10^-1^, *η*^2^_p_=0.003).

**Table 2.** Summary of linear regression results testing the effect of language lateralization phenotype on weighted BOLD asymmetry scores across the five networks supporting visuospatial attention of ALANs. Separate multiple linear regression models were estimated for each network, with weighted BOLD asymmetry scores as the dependent variable. Language lateralization phenotype (typical or atypical) was the primary predictor, controlling for manual preference, age, years of education, total intracranial volume, response hand, and gender. An interaction term between language phenotype and manual preference was also included in all models. The covariates and the interaction term are reported in Table S2. The table reports; the overall model *p*-value (*p*_model_) and its False Discovery Rate (FDR)-corrected value (*p*_FDR_), the adjusted R^2^, the regression coefficient for language phenotype (*β*_Phenotype_), the *t*-value, and associated *p*-values (uncorrected and FDR-corrected), the least-square means with 95% confidence intervals for typical and atypical groups, the group contrast (difference in estimated means) with 95% confidence interval, and the partial eta-squared (*η*^2^_p_) as a measure of effect size. Asterisks indicate networks for which both the main effect of language phenotype and the overall regression model were statistically significant after FDR correction.

| Network | $p_{\text{model}} (p_{\text{FDR}})$ | Adjusted $R^2$ | $\beta_{\text{Phenotype}}$ | $t_{\text{Phenotype}} (df)$ | $p_{\text{Phenotype}} (p_{\text{FDR}})$ | L.S. mean (95% C.I.) | | Contrast (95% C.I.) | $\eta^2_p$ |
| --- | --- | --- | --- | --- | --- | --- | --- | --- | --- |
|  |  |  |  |  |  | Typical | Atypical |  |  |
| Parieto-frontal * | $2.10^{-4}$<br>($4.10^{-4}$) | 0.08 | 0.51 | 2.96 | $3.10^{-3}$<br>($8.10^{-3}$) | 1.73<br>[1.38, 2.08] | 0.72<br>[0.01, 1.43] | 1.01<br>[0.34, 1.69] | 0.031 |
| Temporo-frontal * | $2.10^{-9}$<br>($6.10^{-9}$) | 0.16 | 0.34 | 3.96 | $9.10^{-5}$<br>($5.10^{-4}$) | 0.93<br>[0.76, 1.10] | 0.26<br>[-0.09, 0.61] | 0.67<br>[0.34, 1.00] | 0.054 |
| Posterior-medial | $5.10^{-2}$<br>($5.10^{-2}$) | 0.03 | 0.25 | 2.66 | $1.10^{-2}$<br>($2.10^{-2}$) | 0.95<br>[0.75, 1.15] | 0.44<br>[0.03, 0.85] | 0.51<br>[0.12, 0.90] | 0.023 |
| Somato-motor | $8.10^{-39}$<br>($4.10^{-38}$) | 0.50 | 0.10 | 0.87 | $4.10^{-1}$<br>($4.10^{-1}$) | 0.56<br>[0.33, 0.80] | 0.36<br>[-0.11, 0.84] | 0.20<br>[-0.25, 0.65] | 0.003 |
| Visual * | $8.10^{-4}$<br>($1.10^{-3}$) | 0.07 | 0.27 | 2.42 | $2.10^{-2}$<br>($2.10^{-2}$) | 0.87<br>[0.64, 1.09] | 0.33<br>[-0.12, 0.79] | 0.53<br>[0.10, 0.97] | 0.021 |

**Figure 2.**
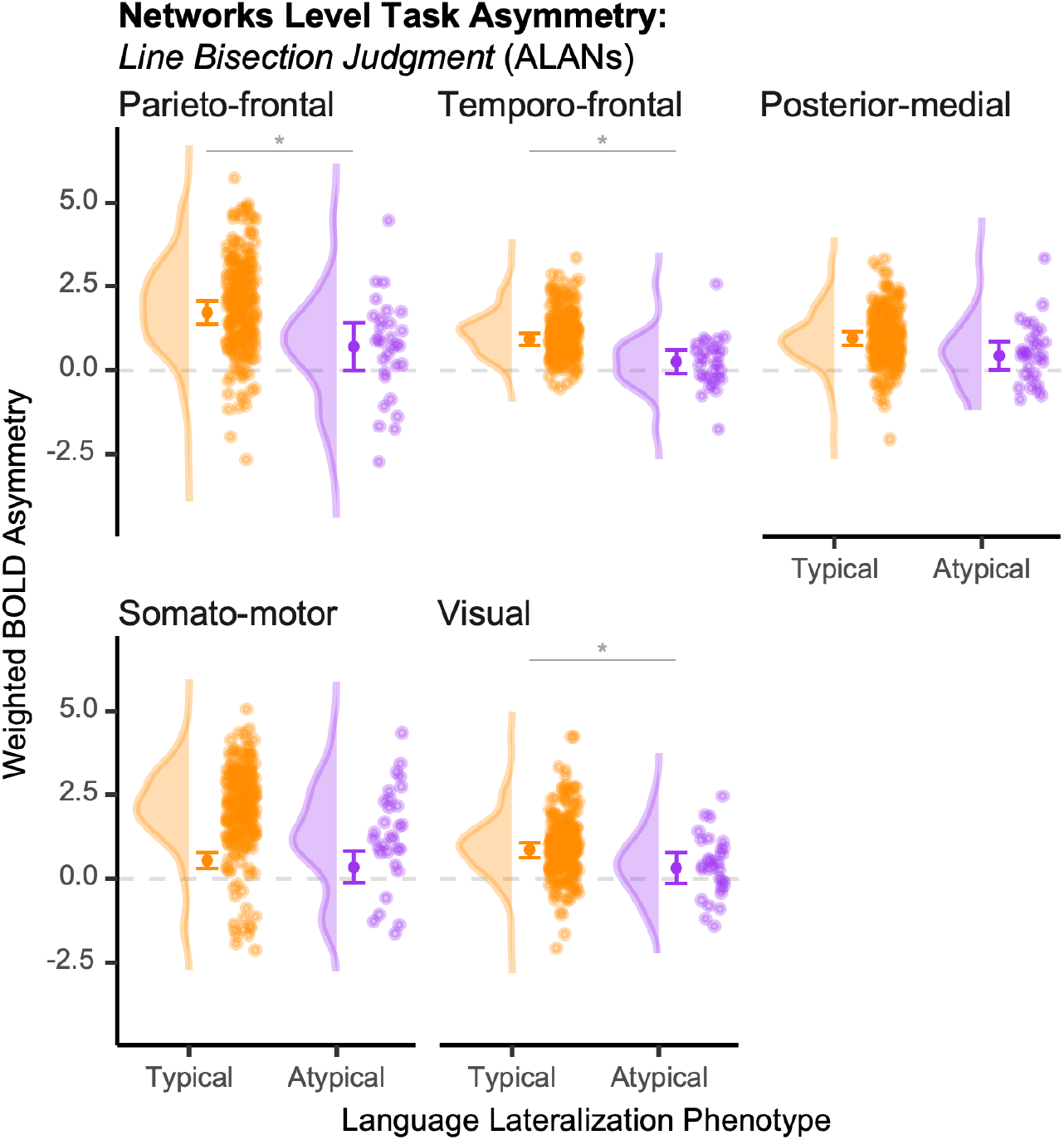
Effect of language lateralization phenotype on weighted BOLD asymmetry scores across the five networks of ALANs. Asymmetry scores are shown for individuals with typical (left-hemisphere dominant; *n*=254, orange) and atypical (right-hemisphere dominant; *n*=30, pink) language lateralization. For each network and group, the figure displays (from left to right): a density plot of the asymmetry score distribution, the estimated marginal mean (central solid dot) with its 95% confidence interval, and a scatter plot of individual asymmetry scores. Asterisks indicate networks for which both the main effect of language lateralization phenotype and the overall regression model were significant after False Discovery Rate correction (*p*_FDR_<0.05, Table 2). For each network, asymmetry was quantified by computing voxel-weighted BOLD activation in the atlas-defined dominant hemisphere versus its homotopic contralateral regions, yielding an asymmetry score (dominant *minus* homotopic network).

Ineach of the threesignificant visuospatial attentional networks, individuals with typical language lateralization exhibited significantly stronger rightward asymmetry scores than atypical individuals during performance of the line bisection judgment task (parieto-frontal: *μ*_Typical_=1.73, C.I._95%_=[1.38, 2.08]; temporo-frontal: *μ*_Typical_=0.93, C.I._95%_=[0.76, 1.10]; visual: *μ*_Typical_=0.87, C.I._95%_=[0.64, 1.09]).

However, the group of atypical individuals did not show reversed (leftward) asymmetries in these networks (Fig. 2). Instead, they displayed bilateral activation patterns in the visual (*μ*_Atypical_=0.33, C.I._95%_=[-0.12, 0.79]) and temporo-frontal (*μ*_Atypical_=0.26, C.I._95%_=[-0.09, 0.61]) networks. In the parieto-frontal network, atypical individuals showed a rightward asymmetry (*μ*_Atypical_=0.72, C.I._95%_=[0.01, 1.43]), yet significantly lower than that of typical individuals (*μ*_Typical_=1.73, C.I._95%_=[1.38, 2.08]).

Importantly, there was no main effect of manual preference, nor a significant interaction between language lateralization phenotype and manual preference (Table S2), indicating that the observed effects were specifically driven by language lateralization phenotype.

Given the bi-hemispheric organization of the temporo-frontal network (18) with 16 rightward lateralized regions and four leftward lateralized regions (Fig. 1a), we further explored whether the effects were specific to one hemisphere. Type III ANCOVA revealed that the observed asymmetry differences were driven primarily by the right-hemisphere component (*p*_model_=1.10^-8^, *p*_Phenotype_=1.10^-4^), whereas the left-hemisphere component, including the Pallidum, the Putamen, the Thalamus and the Superior Temporal Gyrus, did not show a significant phenotype effect (*p*_model_=2.10^-1^, *p*_Phenotype_=3.10^-4^). Atypical individuals displayed a bilateral pattern in the right-hemisphere component (*μ*_Atypical_=0.34, C.I._95%_=[-0.08, 0.77]), whereas typical individuals showed the expected rightward dominance (*μ*_Typical_=1.14, C.I._95%_=[0.94, 1.35]).

### Impact of Language Lateralization Phenotypes on Calculation and Number Interval Comparison Networks Asymmetries

To determine whether language lateralization phenotype also impacts asymmetries in numerical processing networks, we conducted separate linear regression analyses for each of the six networks defined in the Lateralized Underpinnings of Comparison and Arithmetic atlas (LUCA, Fig. 1b, see *Methods*). Analyses were conducted on a subset of 250 BIL&GIN participants (37), using weighted BOLD asymmetry scores as the dependent variable and language lateralization phenotype (typical or atypical) as the primary predictor, while controlling for manual preference, hand response, age, years of education, total intracranial volume, gender, and the interaction between language lateralization phenotype and manual preference.

Type III analysis of covariance revealed a significant main effect of language lateralization phenotype on asymmetry scores in four out of the six networks: both fronto-intraparietal networks associated with the calculation (*p*_model FDR_=1.10^-4^, *p*_Phenotype FDR_=2.10^-4^, *η*^2^ _p_=0.068) and comparison tasks (*p* _model FDR_=5.10^-3^, *p* _Phenotype FDR_=4.10^-2^, *η*^2^ _p_=0.026), the visuo-motor network (*p*_model FDR_=3.10^-2^, *p* _Phenotype FDR_=4.10^-2^, *η*^2^ _p_=0.021) and the sub-cortical network (*p*_model FDR_=3.10^-3^, *p* _*Phenotype FDR*_=4.10^-2^, *η*^2^ _p_=0.022), both recruited during calculation, all surviving FDR correction.

In each of the four significant numerical networks, individuals with typical language lateralization exhibited significantly stronger asymmetry scores than atypical individuals during calculation and comparison tasks (fronto-intraparietal_Calculation_: *μ*_Typical_=0.31, C.I._95%_=[0.24, 0.37]; visuo-motor_Calculation_: *μ*_Typical_=0.18, C.I._95%_=[0.13, 0.23]; sub-cortical_Calculation_: *μ*_Typical_=0.11, C.I._95%_=[0.05, 0.17]; fronto-intraparietal_Comparison_: *μ*_Typical_=0.45, C.I._95%_=[0.27, 0.64]).

Similarly to visuospatial attention processing, the group of atypical individuals did not show reversed asymmetries in these works (Fig. 3). Instead, they displayed bilateral activation patterns (fronto-intraparietal_Calculation_: *μ*_Atypical_=0.05, C.I._95%_=[-0.08, 0.18]; visuo-motor_Calculation_: *μ*_Atypical_=0.08, C.I._95%_=[-0.02, 0.17]; sub-cortical: *μ*_Atypical_=-0.02, C.I._95%_=[-0.14, 0.10]; fronto-intraparietal_Comparison_: *μ*_Atypical_=0.02, C.I._95%_=[-0.34, 0.38]).

**Figure 3.**
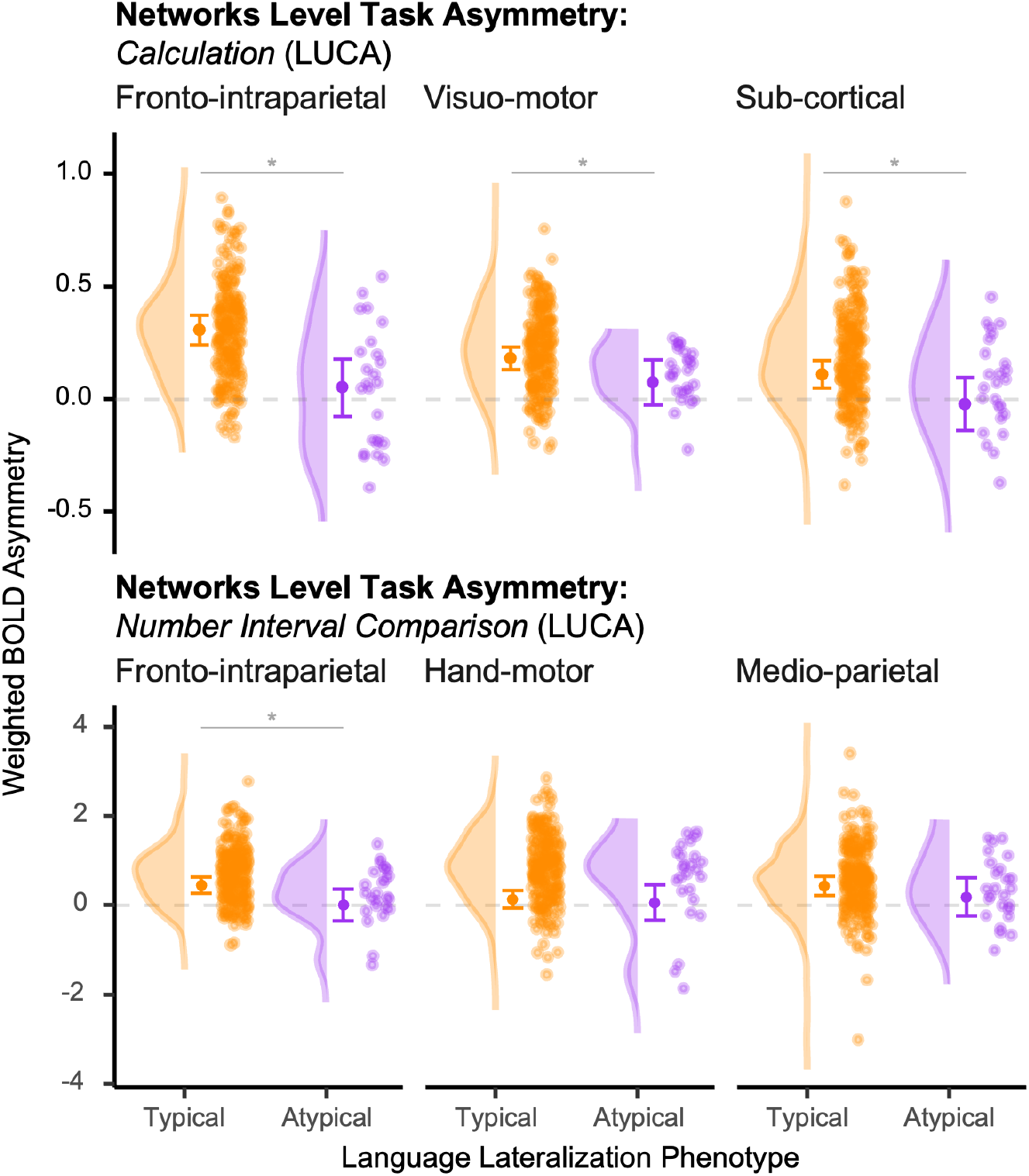
Effect of language lateralization phenotype on weighted BOLD asymmetry scores across the six lateralized networks supporting arithmetics and numerical comparisons of LUCA. Asymmetry scores are shown for individuals with typical (left-hemisphere dominant; *n*=225, orange) and atypical (righthemisphere dominant; *n*=25, pink) language lateralization. For each network and group, the figure displays (from left to right): a density plot of the asymmetry score distribution, the estimated marginal mean (central solid dot) with its 95% confidence interval, and a scatter plot of individual asymmetry scores. Asterisks indicate networks for which both the main effect of language lateralization and the overall regression model were significant after False Discovery Rate correction (*p*_FDR_<0.05, Table 3). For each network, asymmetry was quantified by computing voxel-weighted BOLD activation in the atlas-defined dominant hemisphere versus its homotopic contralateral regions, yielding an asymmetry score (dominant *minus* homotopic network).

As in the visuospatial domain, we found no main effect of manual preference or significant interaction with language phenotype in any of the number networks (Table S3), suggesting that the observed effects were specifically related to hemispheric dominance for language rather than hand preference.

**Table 3.** Summary of linear regression results testing the effect of language lateralization phenotype on weighted BOLD asymmetry scores across the six lateralized networks supporting arithmetics and number interval comparisons of LUCA. Separate multiple linear regression models were estimated for each hub, with weighted BOLD asymmetry scores as the dependent variable. Language lateralization phenotype (typical or atypical) was the primary predictor, controlling for manual preference, age, years of education, total intracranial volume, and gender. An interaction term between language phenotype and manual preference was also included in all models. The covariates and the interaction term are reported in Table S3. The table reports; the overall model *p*-value (*p*_model_) and its False Discovery Rate (FDR)-corrected value (*p*_FDR_), the adjusted R^2^, the regression coefficient for language phenotype (*β*_Phenotype_), the *t*-value, and associated *p*-values (uncorrected and FDR-corrected), the least-square means with 95% confidence intervals for typical and atypical groups, the group contrast (difference in estimated means) with 95% confidence interval, and the partial eta-squared (*η*^2^) as a measure of effect size. Asterisks indicate hubs for which both the main effect of language phenotype and the overall regression model were statistically significant after FDR correction.

| Task | Network | $p_{\text{model}} (p_{\text{FDR}})$ | Adjusted $R^2$ | $\beta_{\text{Phenotype}}$ | $t_{\text{Phenotype}} (df)$ | $p_{\text{Phenotype}} (p_{\text{FDR}})$ | L.S. mean (95% C.I.) | | Contrast (95% C.I.) | $\eta^2_p$ |
| --- | --- | --- | --- | --- | --- | --- | --- | --- | --- | --- |
|  |  |  |  |  |  |  | Typical | Atypical |  |  |
| Calculation | Fronto-intraparietal * | $4.10^{-5}$<br>($1.10^{-4}$) | 0.10 | 0.13 | 4.21 | $4.10^{-5}$<br>($2.10^{-4}$) | 0.31<br>[0.24, 0.37] | 0.05<br>[-0.08, 0.18] | 0.26<br>[0.14, 0.38] | 0.068 |
| | Visuo-motor * | $2.10^{-2}$<br>($3.10^{-2}$) | 0.04 | 0.05 | 2.26 | $2.10^{-2}$<br>($4.10^{-2}$) | 0.18<br>[0.13, 0.23] | 0.08<br>[-0.02, 0.17] | 0.11<br>[0.01, 0.20] | 0.021 |
| | Sub-cortical * | $1.10^{-3}$<br>($3.10^{-3}$) | 0.07 | 0.07 | 2.31 | $2.10^{-2}$<br>($4.10^{-2}$) | 0.11<br>[0.05, 0.17] | -0.02<br>[-0.14, 0.10] | 0.13<br>[0.02, 0.24] | 0.022 |
| Comparison | Fronto-intraparietal * | $3.10^{-3}$<br>($5.10^{-3}$) | 0.06 | 0.22 | 2.53 | $1.10^{-2}$<br>($4.10^{-2}$) | 0.45<br>[0.27, 0.64] | 0.02<br>[-0.34, 0.38] | 0.44<br>[0.10, 0.78] | 0.026 |
| | Hand-motor | $2.10^{-13}$<br>($1.10^{-12}$) | 0.25 | 0.04 | 0.38 | $7.10^{-1}$<br>($7.10^{-1}$) | 0.14<br>[-0.06, 0.34] | 0.07<br>[-0.33, 0.46] | 0.07<br>[-0.30, 0.44] | 0.001 |
| | Medio-parietal | $3.10^{-2}$<br>($3.10^{-2}$) | 0.036 | 0.13 | 1.22 | $2.10^{-1}$<br>($3.10^{-1}$) | 0.44<br>[0.22, 0.66] | 0.19<br>[-0.24, 0.62] | 0.25<br>[-0.15, 0.66] | 0.006 |

Given the bi-hemispheric lateralized organization of the three calculation-related networks (fronto-intraparietal, visuo-motor, and subcortical; see Fig. 1 and Table S1), we further explored whether the effects were specific to one hemisphere. Each network exhibited a distinct asymmetry profile in relation to language lateralization. The effect in the fronto-intraparietal network was driven by its left-hemisphere component (left component: *p*_model FDR_=2.10^-3^, *p*_Phenotype FDR_=5.10^-4^; right component: *p*_model FDR_=7.10^-2^, *p*_Phenotype FDR_=4.10^-2^) with atypical individuals displaying a bilateral pattern in the left component (*μ*_Atypical_=0.02, C.I._95%_=[-0.17, 0.20]), whereas typical individuals showed the expected leftward dominance (*μ*_Typical_=0.35, C.I._95%_=[0.26, 0.45]).

In contrast, the subcortical network showed a significant bilateral effect of language lateralization, driven by its right-hemisphere component (left component: *p*_model FDR_=1.10^-2^, *p*_Phenotype FDR_=2.10^-1^; right component: *p*_model FDR_=6.10^-3^, *p*_Phenotype FDR_=4.10^-2^). Notably, this bilateral network includes only one region in the left hemisphere: the anterior part of the caudate nucleus (Fig. 1, Table S1). Atypical individuals showed a bilateral pattern in the right component (*μ*r_Atypical_=-0.02, C.I.r_95%_=[-0.14, 0.10]), while typical individuals exhibited the expected rightward dominance (*μ*r_Typical_=0.11, C.I.r_95%_=[0.05, 0.17]).

Finally, regarding the visuo-motor network, neither the left nor right component showed a significant effect of language lateralization (left component: *p*r_model FDR_=5.10^-2^, *p*r_Phenotype FDR_=6.10^-2^; right component: *p*r_model FDR_=3.10^-1^, *p*r_Phenotype FDR_=4.10^-1^), suggesting that the observed effect reflects a global bi-hemispheric trend rather than a localized hemispheric difference.

### No Impact of Language Lateralization Phenotype on Intrinsic Markers of Visuospatial Attention and Numerical Networks

Previous research has shown that typical and atypical language lateralization phenotypes are associated with distinct intrinsic connectivity profiles within the language network, including leftward asymmetries in degree centrality, reduced interhemispheric homotopic communication, and lower global integration in typically lateralized individuals (35, 36). To assess whether such differences extend to other lateralized domains, we evaluated the impact of language phenotype on intrinsic connectivity markers in visuospatial attention and calculation and comparison networks.

Similarly to the analyses performed on BOLD task-related asymmetry indices, we conducted separate linear regression analyses for each of the 11 networks, including five from the visuospatial domain (ALANs, Fig. 1) (18) and six from the number processing domain (LUCA, Fig. 1). For each network, we computed three resting-state measures previously associated with language lateralization phenotype (35): degree centrality asymmetry, degree centrality sum, and interhemispheric homotopic intrinsic correlation. All models included language lateralization phenotype (typical or atypical) as the main predictor and controlled for manual preference, age, years of education, total intracranial volume, gender, and response hand.

Type III analysis of covariance revealed no significant main effect of language lateralization phenotype on intrinsic connectivity metrics across either the ALANs or LUCA networks after FDR correction (Tables S4-S9, Fig. S2-S7). All observed effects were small in magnitude (*η*_*2*_^p^≤0.017) with overlapping confidence intervals between gr^2^oups, and no interactions with manual preference were detected.

These results indicate that hemispheric dominance for language does not impact network-level intrinsic connectivity metrics in attentional or numerical atlases, in contrast to our previous findings that it affects intrinsic connectivity metrics within the language network.

## Discussion

Our results indicate that hemispheric functional complementarity does not arise from a single global organizing principle, but instead reflects partially independent lateralization processes shaped by the computational demands of each domain. Importantly, in our framework lateralization is defined using convergent task-evoked and intrinsic connectivity measures for language, visuospatial attention, and numerical cognition. Individuals with typical (left-dominant) language organization; as indexed jointly by task and resting-state metrics, showed the expected pattern of co-lateralized activation for calculation alongside right-lateralized networks for visuospatial attention and numerical comparison. Atypical (right-dominant) individuals, however, did not exhibit a mirror-reversed architecture; rather, they displayed systematically reduced asymmetry and greater bilaterality across both language-aligned and complementary functions. Notably, these task-evoked differences were not mirrored in intrinsic connectivity, as resting-state measures revealed no systematic effect of language phenotype within attentional or numerical networks. Taken together, these findings argue against strong reciprocal models of cross-domain lateralization and instead support a graded-coupling framework in which functional asymmetries can covary but are not obligatorily linked.

Previous work has suggested that interhemispheric connectivity, white-matter architecture, and developmental timing constrain the emergence of lateralization patterns (40–42). In particular, atypical language organization often reflects bilateral recruitment or a shift in the balance of interhemispheric cooperation rather than a simple inversion of the canonical pattern (15, 23, 35). This view aligns with our observation that atypical individuals for language do not exhibit a complete reversal of attentional or numerical asymmetries. Notably, large-sample syntheses emphasize that any cognitive advantages associated with the “typical” (left hemisphere language/right hemisphere visuospatial) pattern are modest at best and that the strength or bilaterality of lateralization may be more predictive of behavior than its direction per se, emphasizing our focus on asymmetry magnitude across domains (43). Furthermore, although language lateralization appears to exert broad effects on large-scale information integration (36), this influence does not appear to translate into local network connectivity changes within non-language networks. In other words, a language lateralization “signature” may be evident in global brain organization without perturbing the fine-grained connectivity of other domain-specific networks. Moreover, this apparent null effect on resting-state connectivity is observed in healthy subjects (*i*.*e*., the language phenotype had no detectable effect on non-language network resting connectivity in our sample). By contrast in patient populations, language-related pathology might induce persistent reorganization even at rest, potentially ‘fixing’ alterations in non-language networks (44, 45), but in our healthy atypical subjects no such effect is seen. These observations strengthen our interpretation that language lateralization exerts domain-global influences rather than inducing local network rewiring.

Number and calculation tasks engage distinct lateralization profiles: calculation tends to recruit a left-lateralized parietal-frontal circuit, whereas numerical interval comparison engages a more bilateral or right-leaning parietal network (46). Visuospatial attention networks, particularly those anchored in the right inferior parietal cortex, retain their lateralization across individuals (30, 34), even when language dominance varies. These findings align with a modular organization in which each domain optimizes its neural configuration according to task demands and evolutionary pressures (47, 48). Extending this modular view to language-attention interplay, new evidence localizes a frontal-eye-field hub where reading and attention interact, and shows that its structural communicability links dorsal attention and sublexical/oculomotor circuits (49). This provides a mechanistic substrate for task-specific co-recruitment without implying global cross-domain coupling.

In fact, one can conceptualize a hierarchical gradient ranging from highly domain-specific organization during task performance, to organization during tasks of other domains, and finally to cortex-wide gradients observable at rest, where purely local resting-state markers show no detectable effect. This hierarchy suggests that the influence of lateralization “cascades” from strong domain-specific task activation, and ultimately to large-scale resting-state architecture, with diminishing magnitude along this cascade. Further, hemispheric interaction asymmetries may help explain why language lateralization does not necessarily drive attention lateralization: left-hemisphere regions are biased to interact more strongly within the same hemisphere, whereas right-hemisphere regions exhibit stronger interhemispheric interactions (50).

From a systems perspective, functional lateralization reflects the interplay between specialized processing hubs and integrative network cores. Yeo and colleagues showed that the association cortex exhibits a mosaic of regions, some with stable functional affiliations and others with flexible connectivity patterns (51). Our resting-state results, showing no systematic differences in non-language network connectivity across language phenotypes, suggest that these flexible hubs preserve their integrative roles irrespective of language dominance. This is compatible with the proposals that domain-specific networks operate within a partially shared structural and energetic scaffold (23, 50) without enforcing rigid cross-domain coupling. Converging with this, the frontal-eye-field reading interaction described above situates a control/interface node within the dorsal attention network rather than prompting wholesale system-wide reconfiguration, again favoring local over global coupling (49).

Critically, cross-domain complementarity can emerge under specific trait constraints. Indeed, the association between language and spatial attention lateralization has been shown to depend on manual preference strength: strong left-handers exhibit a negative correlation (more right-hemisphere spatial with more left-hemisphere language), whereas right- and mixed-handers do not, consistent with largely independent lateralization in other domains (39). This pattern aligns with our findings of stronger asymmetries in typical individuals, while atypical individuals show no mirror reversal.

Taken together, these findings argue against a universal reciprocal mechanism in which dominance for one function forces complementary dominance in another. Instead, they point to a hybrid model: cross-domain correlations arise from shared anatomical constraints, interface hubs, and overlapping developmental windows, but each function retains substantial independence in how, when, and whether it lateralizes. This interpretation reconciles elements of both the causal and statistical hypotheses, while also aligning with proposals that lateralization reflects an adaptive balance between modular specialization, interhemispheric integration, and domain-general control. The graded-coupling framework accommodates trait-conditional coupling and task-conditional co-recruitment without invoking a single global driver (39, 43, 49).

Taken together, these findings redefine our understanding of hemispheric specialization. Rather than reflecting a single global principle linking all lateralized functions, the human brain emerges as a mosaic of domain-specific asymmetries, each optimized to its computational demands yet coordinated within a flexible, integrative scaffold. Typical left-hemisphere language dominance is linked to amplified lateralization across attention and numerical systems, but its atypical variant does not invert this organization; revealing that “reversed” brains are not mirror images but differently balanced systems. The dissociation between extrinsic (task-evoked) and intrinsic (resting-state) organization highlights a hierarchical aspect of brain architecture: lateralization is dynamically expressed under cognition demand, while remaining sufficiently stable to maintain interhemispheric balance at res. By integrating multimodal, multi-domain imaging in a uniquely large phenotyped cohort, this work challenges canonical models of hemispheric complementarity and supports a graded-coupling framework in which specialization is not a fixed trait but a flexible, adaptive strategy of the human brain.

## Methods

### Participants

The study sample consisted in a total of 287 participants from the BIL&GIN (38) whose brain lateralization for language has been previously identified as either (35): typical (left hemisphere dominant for language; *n*=257, 125 left-handers, 125 women) or atypical (right hemisphere dominant for language; *n*=30, 25 left-handers, 15 women). The mean age of the sample was 25.8 years (*σ*=6.5; range: 18 − 57 years), and the mean level of education was 15.6 years (*σ*=2.3 years; range: 11 − 20 years), corresponding to almost five years of education after the French baccalaureate.

All participants were free of brain abnormalities as assessed by a trained radiologist inspecting their structural T1-MRI scans. All participants gave their informed written consent and received compensation for their participation. The Basse-Normandie Ethics Committee approved the study protocol.

Depending on the analysis, different subsamples of participants completed resting-state fMRI and/or task-based fMRI sessions (Table 1), including a visuospatial session with a line bisection judgment task and a numerical session comprising calculation and number interval comparison tasks (38).

### Behavioral Tasks

Visuospatial attention was evaluated using a line bisection judgment task described in the BIL&GIN data release paper (38). On each trial, participants (*n*=284) viewed a horizontal line bisected by a short vertical segment (subtending 1° of visual angle) for 2 seconds, followed by a 10-second delay during which only a central fixation cross was displayed. Participants indicated whether the bisection mark was centered, shifted to the left, or shifted to the right relative to the true midpoint, using a three-button response pad operated preferentially with the right hand (index finger for “left,” middle finger for “center,” and ring finger for “right”). Stimuli varied in horizontal position (−7°, 0°, or +7° offset) and line length (6°, 7°, or 9° of visual angle), with the bisection mark offset by 0.3° to the left or right. Conditions were fully counterbalanced. Participants completed 36 trials, equally distributed across centered, leftward-, and rightward-bisected lines. A 12-second fixation period was presented before the first trial and after the final trial. Prior to scanning, participants completed a practice session to ensure task comprehension.

Numerical cognition was assessed using two tasks, fully described in the BIL&GIN data release paper (38): a calculation task and a number interval comparison task. In the calculation task, participants (*n*=250) mentally added three two-digit numbers (*e*.*g*., 25+12+4). Each trial lasted up to 8 seconds, during which participants pressed a button as soon as they had computed the result in their mind. Thus, no in-scanner performance accuracy was collected. Task performance was assessed immediately after scanning during a debriefing session, in which participants completed the same run again and reported their answers aloud. Each trial was followed by a 6-second baseline during which participants monitored a central fixation cross for subtle visual changes.

In the number interval comparison task, participants viewed a triplet of two-digit numbers and indicated which interval was numerically larger; the one between the center and left number (*e*.*g*., 31–56) or the one between the center and right number (*e*.*g*., 56–95). Responses were made using a two-button response pad operated with the right hand: left button for a larger left interval, right button for a larger right interval. Each trial lasted 5 seconds, followed by a 7-second fixation baseline. For both tasks, each run began and ended with a short fixation period, and participants completed practice trials prior to scanning to ensure comprehension of task instructions.

### MRI Acquisition and Preprocessing

Structural and functional MRI data were acquired on a 3T Philips Intera Achieva scanner (Philips, Eindhoven, The Netherlands), following procedures described by Mazoyer and colleagues (38).

#### Structural Imaging

High-resolution T1-weighted images (T1w) were obtained using a 3D turbo field echo sequence (TR=20 ms; TE=4.6 ms; flip angle=10°; inversion time=800 ms; turbo factor=65; SENSE factor=2; field of view=256×256×180 mm^3^; voxel size=1×1×1 mm^3^). The acquisition plane was aligned along the anterior-posterior commissural line. In addition, T2*-weighted images were collected using a fast field echo sequence (TR=3,500 ms; TE=35 ms; flip angle=90°; SENSE factor=2; 70 axial slices; voxel size=2×2×2 mm^3^).

#### Functional Imaging

Task-related fMRI data were acquired using a T2*-weighted echo-planar imaging sequence (TR=2 s; TE=35 ms; flip angle=80°; 31 axial slices; field of view=240×240 mm^2^; voxel size=3.75×3.75×3.75 mm^3^). The first four volumes were discarded to allow MR signal stabilization.

### Preprocessing and Analysis

Data preprocessing was performed using SPM12 (www.fil.ion.ucl.ac.uk/spm/) with custom MATLAB scripts. T2*-FFE images were coregistered to the T1w anatomical scans, which were segmented and normalized to the BIL&GIN template (aligned to MNI space). Functional data underwent slice timing and motion correction; motion parameters were regressed out. T2*-EPI volumes were normalized to standard space (2×2×2 mm^3^) using combined transformations and trilinear interpolation.

For resting-state fMRI data, additional nuisance regressors were removed, including average signals from white matter and cerebrospinal fluid compartments, along with linear temporal trends. Time series were then bandpass filtered (0.01–0.1 Hz) using a least-squares linear-phase finite impulse response filter. Individual regional BOLD time series were computed by averaging the signal across all voxels within each region of interest.

Task-related fMRI responses were analyzed with a general linear model in SPM12. Functional volumes were smoothed with a 6 mm FWHM Gaussian kernel and high-pass filtered (159 s). Trial events were modeled as boxcar functions (2 s duration) convolved with a canonical hemodynamic response function. Contrast maps were generated at the individual level and subjected to region-of-interest analysis using the AICHA atlas (52). BOLD signal changes were extracted from 185 pairs of homotopic regions, excluding seven pairs with susceptibility artifacts.

The AICHA atlas (52) was chosen because it accounts for hemispheric torsion (Yakovlevian torque (1)) and allows reliable estimation of functional asymmetries in homologous cortical regions.

### Preprocessing and Analysis

Statistical analyses were conducted using R (53) (R version: 4.5.0). Data wrangling, statistics and visualization were performed using the R libraries *car* (54) (R package version: 3.1-3), *dplyr* (55) (R package version: 1.1.4), *tidyr* (R package version: 1.3.1), *purrr* (57) (R package version: 1.0.4), *broom* (58) (R package version: 1.0.8), *emmeans* (59) (R package version: 1.11.0), effectsize (60) (R package version: 1.0.0), and *ggplot2* (61) (R package version: 3.5.2). Brain visualizations were realized using Surf Ice (62, 63), and were made reproducible following guidelines to generate programmatic neuroimaging visualizations (64).

### Visuospatial Task-Based Network Asymmetries Analyses

To assess the impact of language lateralization on visuospatial attention networks organization during the line bisection judgment task, separate linear regression models were estimated for each of the five networks of the Atlas for the Lateralized visuospatial Attention Networks (ALANs (18), Fig. 1): parieto-frontal, temporo-frontal, posterior-medial, somato-motor, and visual. For each network, a weighted BOLD asymmetry score was computed to quantify task-related lateralization. Specifically, for each region within the hemisphere originally identified as dominant for a given network (as defined in ALANs), activation values were weighted by the number of voxels in that region. These values were summed across all dominant-hemisphere regions and divided by the total voxel count of the network. The same procedure was applied to homotopic regions in the contralateral hemisphere. Asymmetry scores were then computed as: atlas-defined network *minus* homotopic network.

Language lateralization phenotype (typical or atypical, as defined by Labache and colleagues (65) was the primary predictor, with manual preference included as an additional factor. Covariates of no interest included age, years of education, total intracranial volume, gender, and response hand). An interaction term between language lateralization phenotype and manual preference was included to assess combined effects.

Given the bilateral nature of the Temporo-frontal network (18) with 16 regions in the right hemisphere and four in the left (Fig. 1), we conducted additional analyses examining asymmetry separately within left and right sub-regions using the same regression and post hoc procedures. Model significance was evaluated using the False Discovery Rate (66) procedure, denoted as *p*r_FDR_. For models reaching significance, post hoc analyses were conducted using Type III sum-of-squares analysis of variance to assess individual predictors. The specific effect of language phenotype was examined using estimated marginal means (least-squares means), adjusted for all covariates. Pairwise contrasts were tested using Tukey’s HSD.

#### Extension to Number Tasks

To further investigate hemispheric functional complementarity, we extended our approach to two distinct numerical cognition tasks (38), previously shown to elicit lateralized patterns of activation: an arithmetic calculation task and a numerical interval comparison task. These tasks are known to differentially engage hemispheric networks, with arithmetic operations typically recruiting predominantly left-lateralized regions, while numerical interval comparisons preferentially activate right-hemispheric circuits.

Following previously published methods for constructing cognitive atlases of lateralized function (16–18), we identified the anatomo-functional basis of these two tasks using a reference group of 120 right-handed participants with typical language lateralization (35). We used a conjunction analysis to define relevant regions for both the calculation and numerical comparison task: 1) regions showing significantly leftward activation and asymmetry (threshold: *p*=(0.05/185)^2^=7.10^-8^, Bonferroni-corrected for 185 regions), and 2) regions showing significantly rightward activation and asymmetry at the same threshold. This resulted in a total of 60 brain regions. The calculation task recruited 43 brain regions; 23 in the left hemisphere, and 20 in the right. The comparison task recruited 37 brain regions; 9 in the left hemisphere, and 28 in the right. Resting-state connectivity analyses were then conducted separately for each task-specific set of regions. The regions engaged during calculation were grouped into three networks (Fig. 1, Table S1): a fronto-intraparietal network spanning superior, middle, and inferior frontal gyri through the intraparietal sulcus, supplementary motor area, anterior cingulate, and orbitofrontal cortex, supporting symbolic manipulation; a visuo-motor network linking lingual, fusiform, inferior occipital, and inferior temporal regions with pre- and postcentral cortices and the Rolandic operculum, integrating visual numeral processing with sensorimotor coding and subvocal articulation; and a subcortical network comprising thalamus, caudate, hippocampus, amygdala, and parahippocampal gyrus, contributing gating, memory, and arousal modulation. Comparison-related activity was also organized into three distinct networks (Fig. 1, Table S1): a fronto-intraparietal system extending from frontal to intraparietal regions mediating attentional selection and evidence-based decisions; a hand-motor response network encompassing posterior occipito-temporal and peri-Rolandic regions, supporting visual numeral analysis and response preparation; and a medio-parietal network centered on the precuneus and parieto-occipital sulcus, emphasizing midline parietal hubs for internal evidence accumulation and spatial-numeric mapping. This parcellation defines the Lateralized Underpinnings of Comparison and Arithmetic atlas (LUCA; Fig. 1, Table S1).

We then computed weighted BOLD asymmetry scores (atlas-defined network *minus* homotopic network) for each of the six LUCA networks and assessed the effect of language lateralization phenotype using the same linear modeling framework as described above.

### Intrinsic Connectivity Analyses

Finally, building on the revised definitions of typical and atypical language phenotypes (36, 65), we assessed whether language lateralization also affects intrinsic markers of lateralised networks organization in both visuospatial attention (ALANs) and numerical cognition (LUCA). For each of the 11 networks in total (five from ALANs and six from LUCA, Fig. 1), we computed three intrinsic connectivity metrics: degree centrality asymmetry, degree centrality sum, and interhemispheric homotopic intrinsic correlation, as defined by Labache and colleagues (35).

Briefly, degree centrality was computed for each region as the sum of its positive correlations with all other regions within the same network, indexing regional integration. For each network, degree centrality values were then averaged separately within the atlas-defined hemisphere and its homotopic counterpart. The degree centrality asymmetry score was calculated as: atlas-defined network minus homotopic network. The degree centrality sum was calculated as: (atlas-defined network + homotopic network)/2.

To evaluate the influence of language lateralization phenotype on these intrinsic connectivity metrics, we applied the same covariate-adjusted linear modeling framework used for task-based BOLD asymmetry analyses.

## Acknowledgments

The results are part of the BIL&GIN database (38).

## Data and Code Availability Statement

The data and the code used in the Method section to process the data, to reproduce the results and visualizations can be found here: https://github.com/loiclabache/Labache_2025_IndLat. The atlas for the lateralized visuospatial attention networks (ALANs) can be found here (67): https://github.com/loiclabache/ALANs_brainAtlas. The lateralized underpinnings of comparison and arithmetic atlas can be found here: https://github.com/loiclabache/LUCA_brainAtlas.

## CRediT Authorship Contribution Statement

**Loïc Labache:** Conceptualization, Data Curation, Formal Analysis, Investigation, Methodology, Project administration, Resources, Supervision, Software, Validation, Visualization, Writing—original draft, Writing—review & editing, Supervision, and Project administration. **Isabelle Hesling:** Investigation, Project administration, Resources, Validation, Writing— original draft, Writing—review & editing. **Laure Zago:** Conceptualization, Data Curation, Funding acquisition, Investigation, Methodology, Resources, Validation, Visualization, Writing—original draft, Writing—review & editing.

## Extended Data Figures

**Extended Data Fig. 1.**
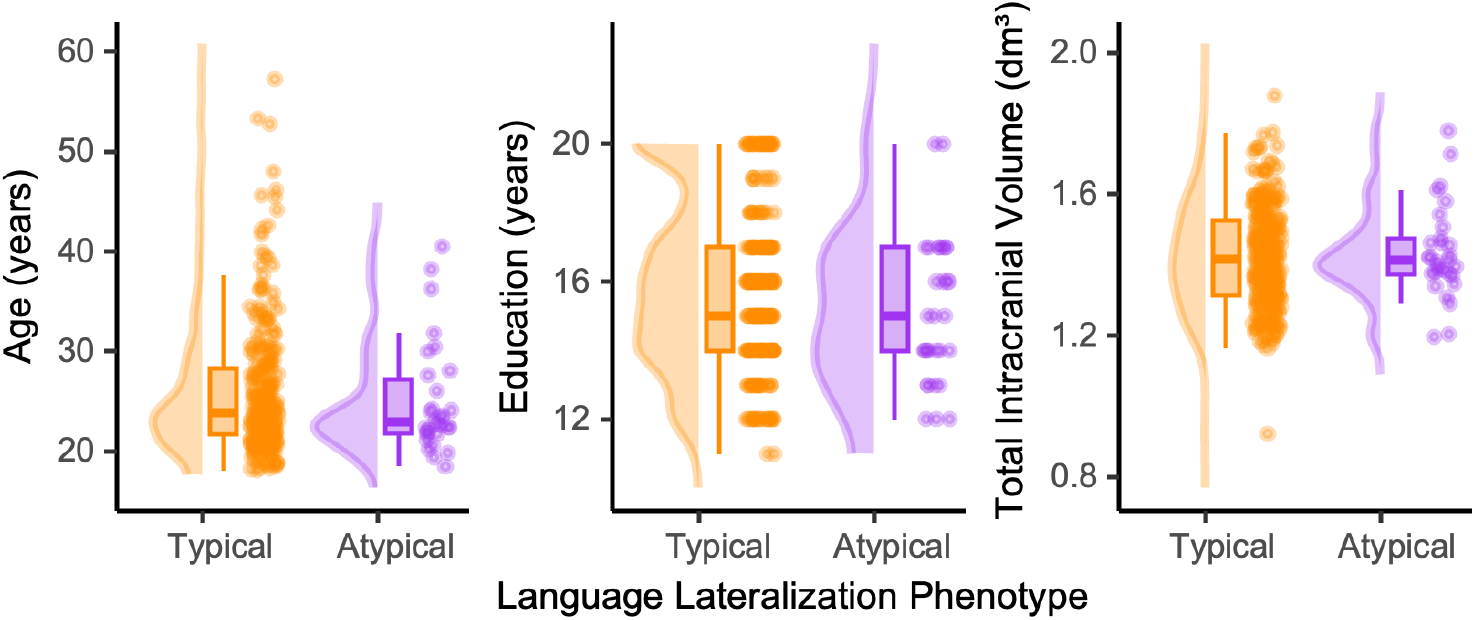
Distribution of covariates across language lateralization phenotypes. Each plot illustrates the distributions of age (left), years of education (middle), and total intracranial volume (right) for individuals with typical (left-hemisphere dominant; orange) and atypical (right-hemisphere dominant; purple) language lateralization. For each covariate and group, the figure displays (from left to right): a density plot of the covariate distribution, a boxplot representing the median and interquartile range, and a scatter plot of individual values. These covariates were included in all regression models (see Table S2).

**Extended Data Fig. 2.**
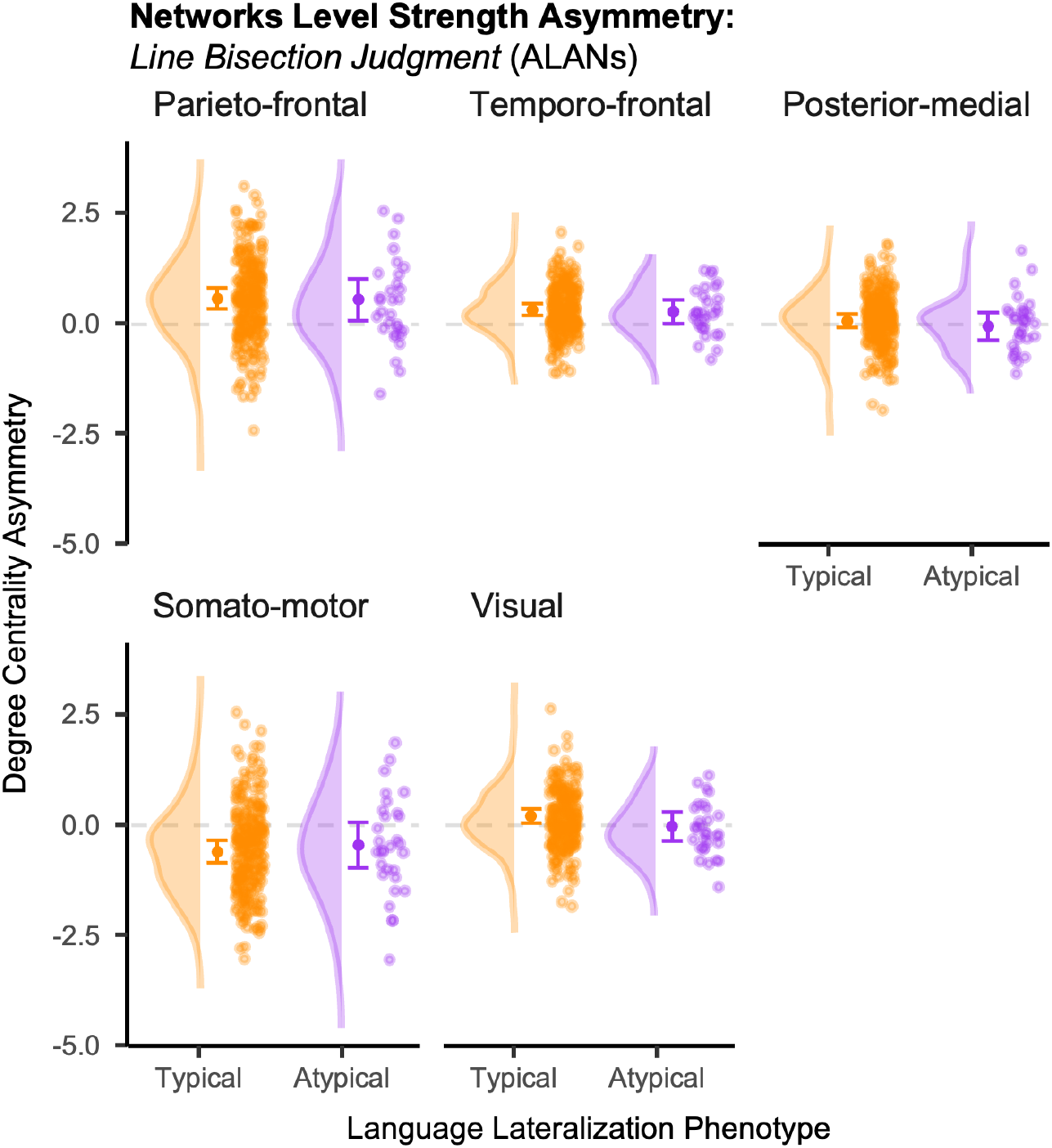
Effect of language lateralization phenotype on the degree centrality asymmetry across the five visuospatial attention networks. Degree centralities are shown for individuals with typical (left-hemisphere dominant; *n*=254, orange) and atypical (right-hemisphere dominant; *n*=30, pink) language lateralization. For each network and group, the figure displays (from left to right): a density plot of the correlation distribution, the estimated marginal mean (central solid dot) with its 95% confidence interval (Table S4), and a scatter plot of individual degree centralities.

**Extended Data Fig. 3.**
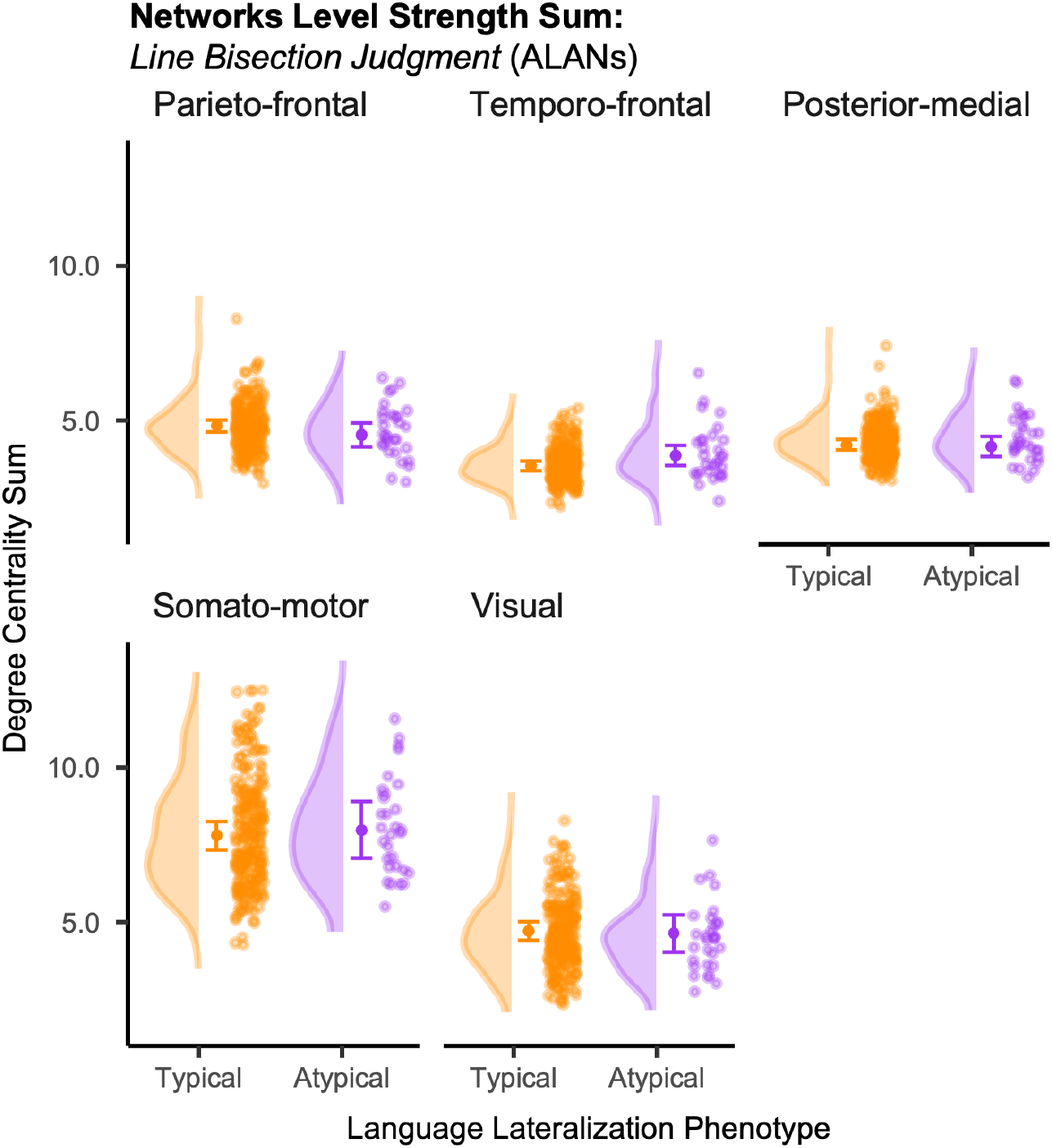
Effect of language lateralization phenotype on the degree centrality sum across the five visuospatial attention networks. Degree centralities are shown for individuals with typical (left-hemisphere dominant; *n*=254, orange) and atypical (right-hemisphere dominant; *n*=30, pink) language lateralization. For each network and group, the figure displays (from left to right): a density plot of the correlation distribution, the estimated marginal mean (central solid dot) with its 95% confidence interval (Table S5), and a scatter plot of individual degree centralities.

**Extended Data Fig. 4.**
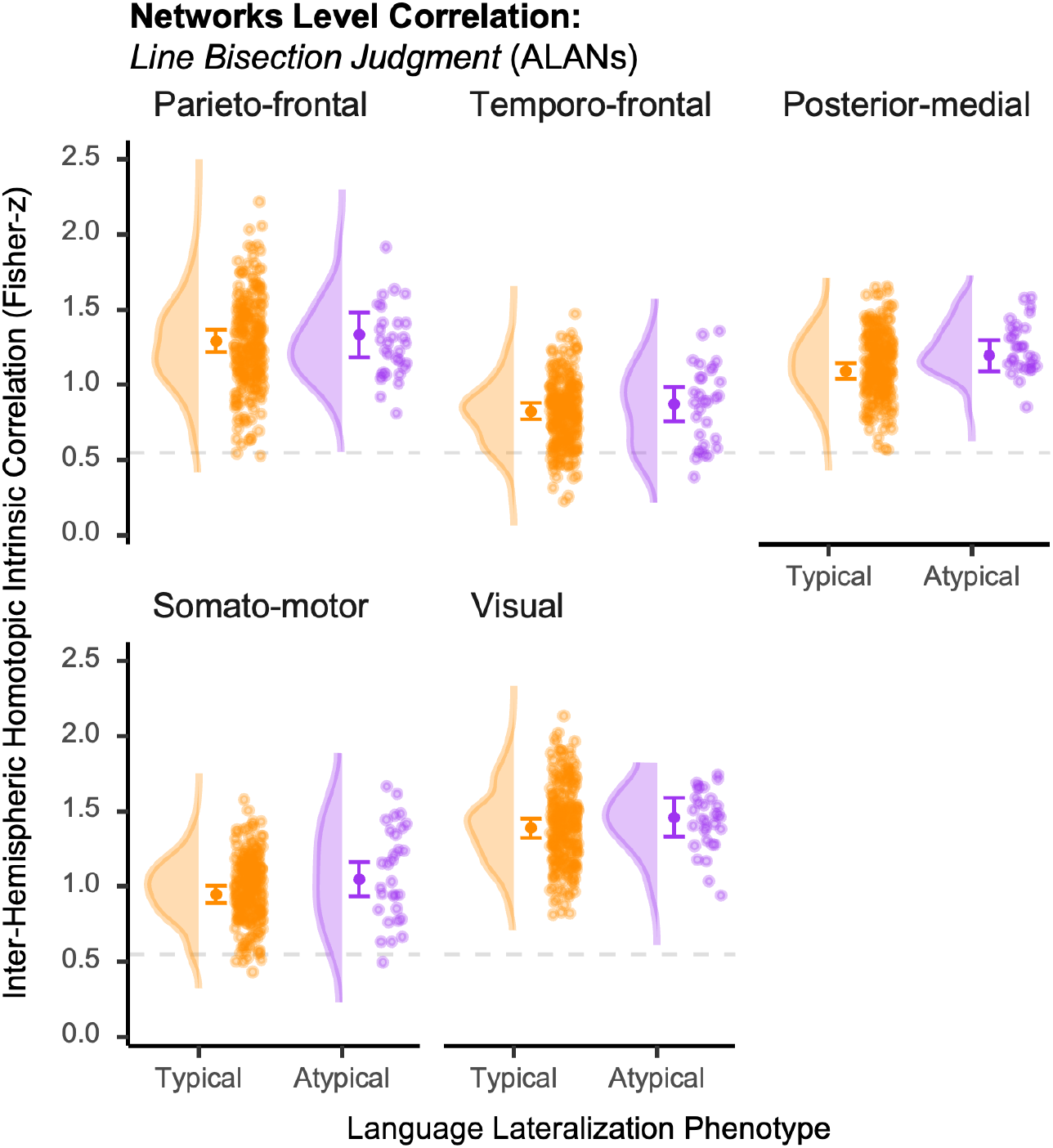
Effect of language lateralization phenotype on the inter-hemispheric homotopic intrinsic correlation (Fisher *z*-transformed) across the five visuospatial attention networks. Correlations are shown for individuals with typical (left-hemisphere dominant; *n*=254, orange) and atypical (right-hemisphere dominant; *n*=30, pink) language lateralization. For each network and group, the figure displays (from left to right): a density plot of the correlation distribution, the estimated marginal mean (central solid dot) with its 95% confidence interval (Table S6), and a scatter plot of individual correlations.

**Extended Data Fig. 5.**
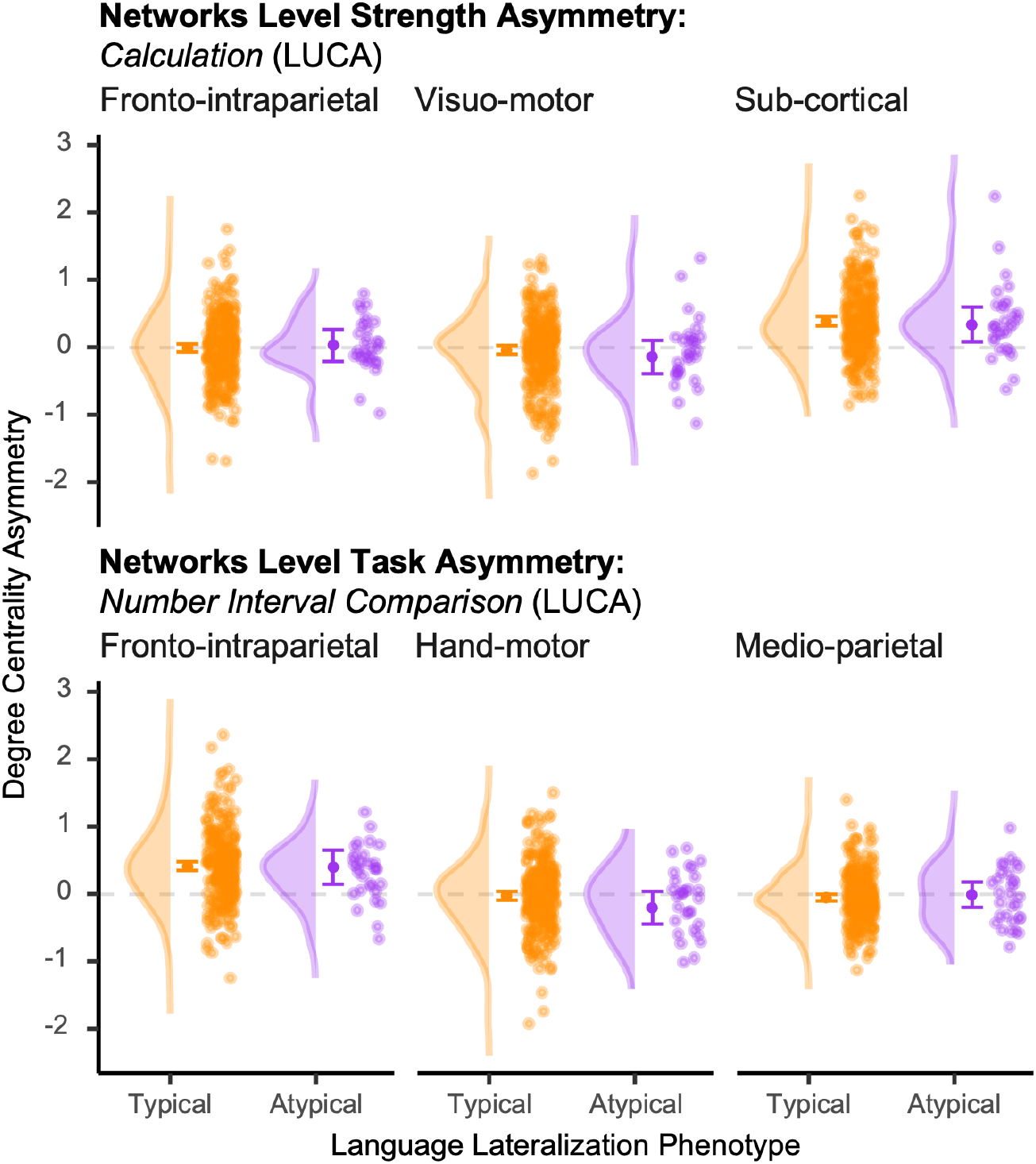
Effect of language lateralization phenotype on the degree of centrality asymmetry across the six lateralized networks supporting arithmetics and numerical comparisons of LUCA. Degree centralities are shown for individuals with typical (left-hemisphere dominant; *n*=257, orange) and atypical (right-hemisphere dominant; *n*=30, pink) language lateralization. For each network and group, the figure displays (from left to right): a density plot of the correlation distribution, the estimated marginal mean (central solid dot) with its 95% confidence interval (Table S7), and a scatter plot of individual degree centralities.

**Extended Data Fig. 6.**
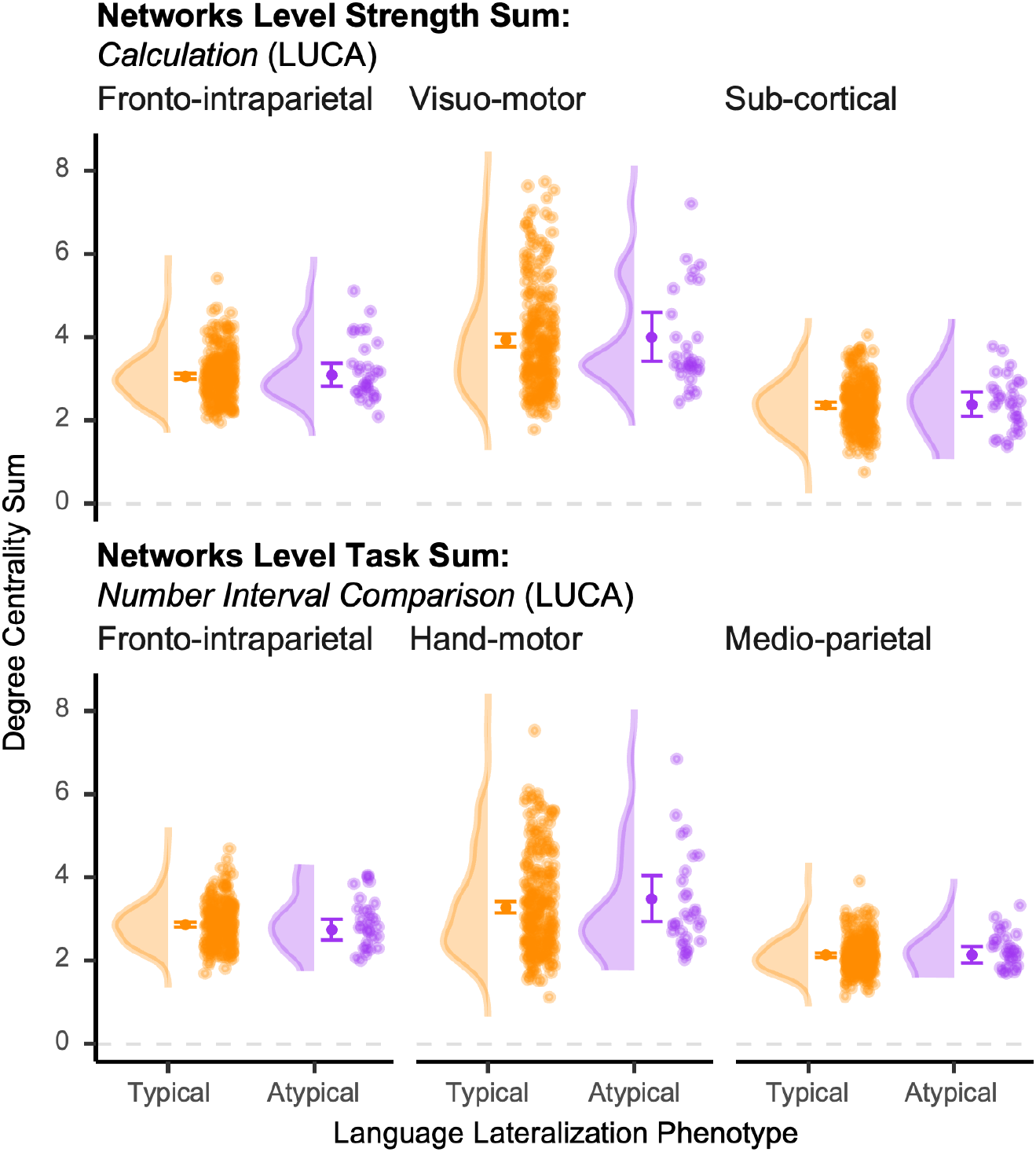
Effect of language lateralization phenotype on the degree centrality sum across the six lateralized networks supporting arithmetics and numerical comparisons of LUCA. Degree centralities are shown for individuals with typical (left-hemisphere dominant; *n*=257, orange) and atypical (right-hemisphere dominant; *n*=30, pink) language lateralization. For each network and group, the figure displays (from left to right): a density plot of the correlation distribution, the estimated marginal mean (central solid dot) with its 95% confidence interval (Table S8), and a scatter plot of individual degree centralities.

**Extended Data Fig. 7.**
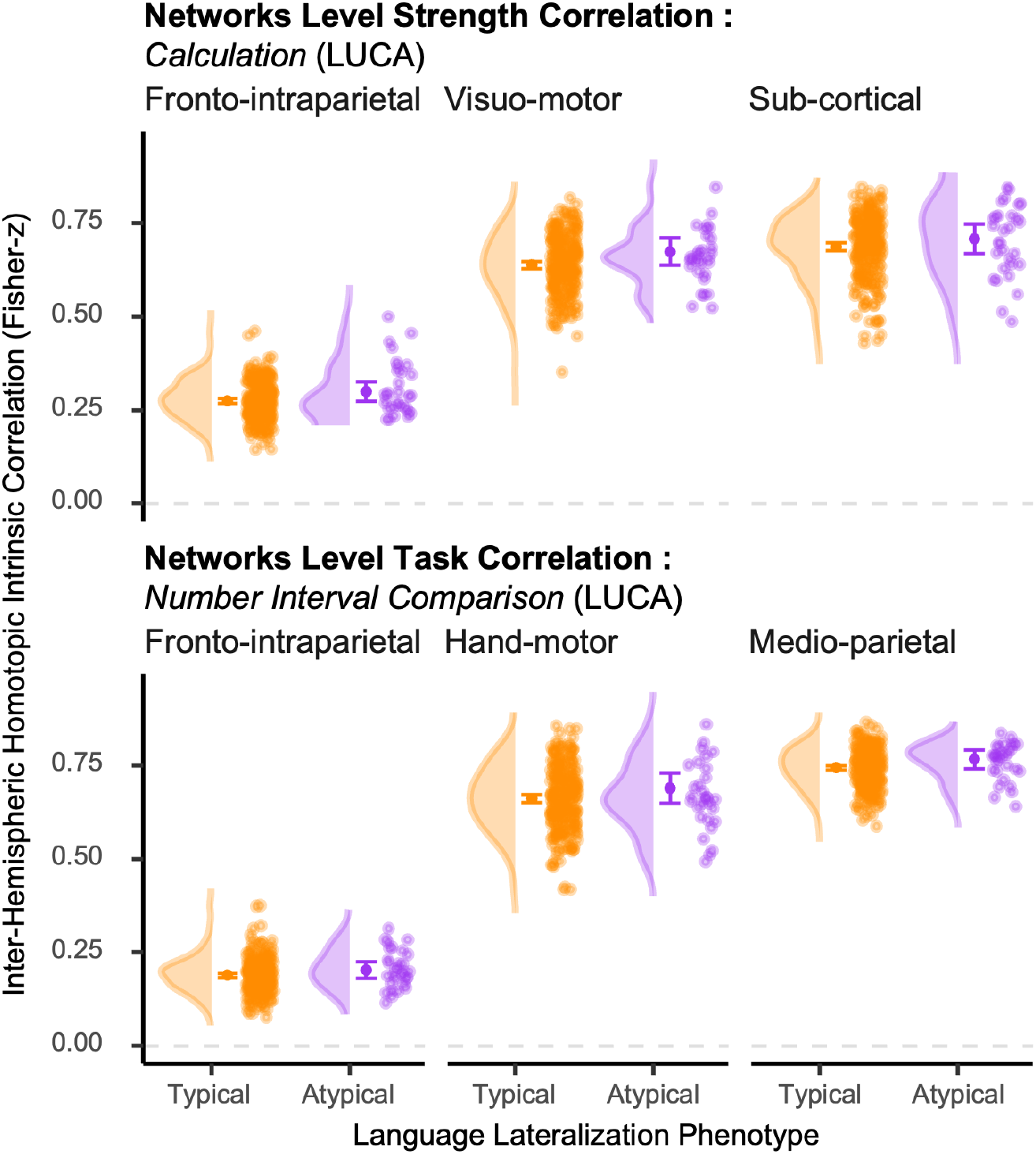
Effect of language lateralization phenotype on the inter-hemispheric homotopic intrinsic correlation (Fisher *z*-transformed) across the six lateralized networks supporting arithmetics and numerical comparisons of LUCA. Correlations are shown for individuals with typical (left-hemisphere dominant; *n*=257, orange) and atypical (right-hemisphere dominant; *n*=30, pink) language lateralization. For each network and group, the figure displays (from left to right): a density plot of the correlation distribution, the estimated marginal mean (central solid dot) with its 95% confidence interval (Table S9), and a scatter plot of individual correlations.

## Extended Data Tables

**Extended Data Table 1.**
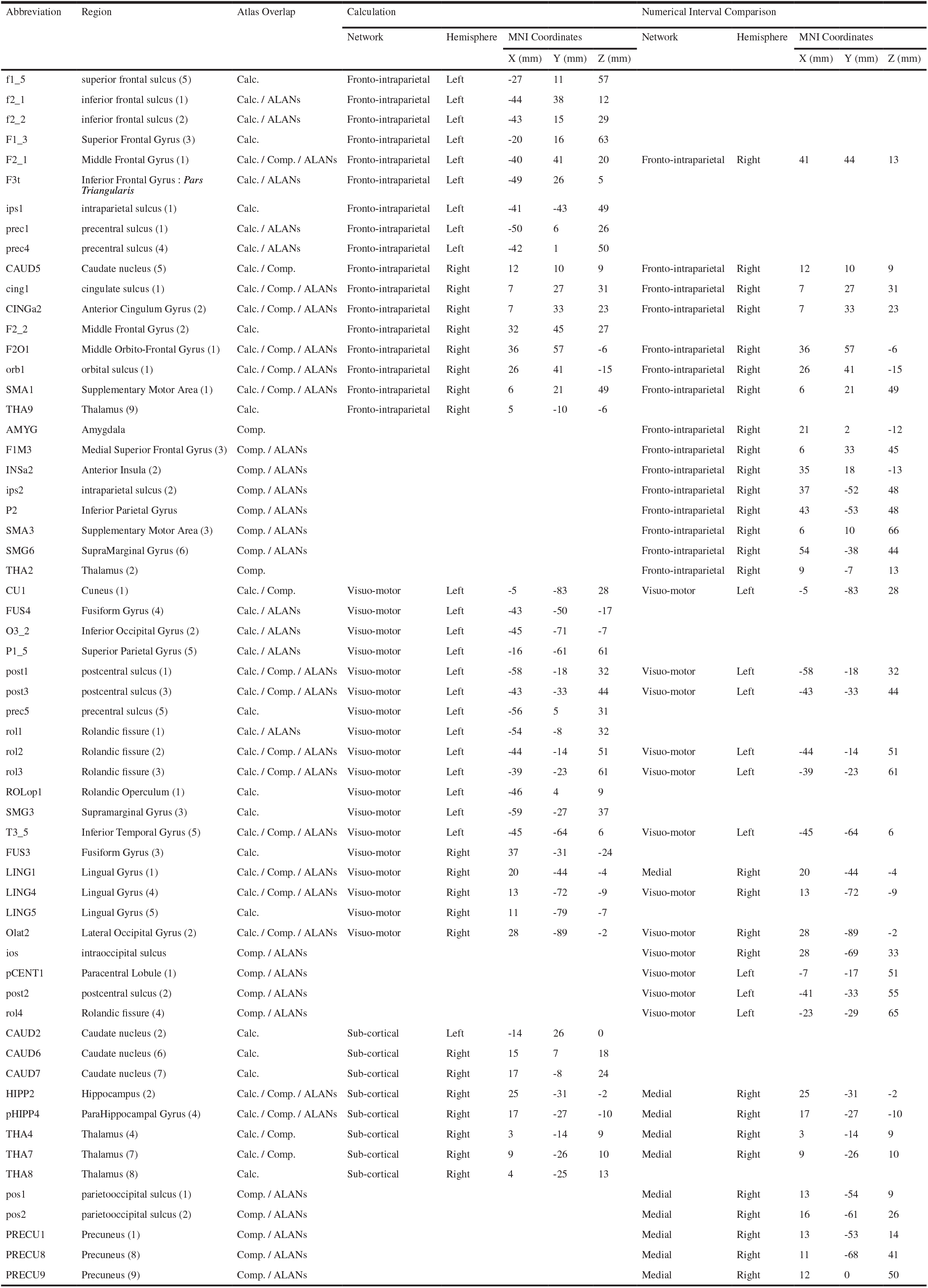
Description of the Lateralized Underpinnings of Comparison and Arithmetic atlas (LUCA, Fig. 1). For both tasks included in LUCA (*Calculation* and *Numerical Interval Comparison*): label of the network to which a region has been clustered (*Network*), its abbreviation (*Abbreviation*), its full anatomical name (*Region*), the atlas label(s) in which the region is represented, determined from the Comparison, Calculation, and ALANs parcellations (*Atlas Overlap*; multiple atlas names indicate regions present in more than one parcellation; *Calc*., Calculation; *Comp*., Comparison; *ALANs*, Atlas for Lateralized visuospatial Attentional Networks (18)), the hemisphere to which it belongs (*Hemisphere*), and the coordinates of its center of mass in MNI space (*MNI Coordinates*: *X, Y*, and *Z*). The number in parentheses in the Region column corresponds to the functional subdivision of the region. The names of the regions correspond to those defined in the AICHA atlas (52).

**Extended Data Table 2.** Full regression results for covariates and interaction term in models assessing the effect of language lateralization phenotype on weighted BOLD asymmetry scores across the five visuospatial attention networks. This table reports the full results for all covariates and the interaction term (language phenotype by manual preference) from the linear regression models presented in Table 2. For each of the five visuospatial attention networks, the following are shown for each predictor: regression coefficient (*β*_Predictor_), standard error (S.E.), *t*-value, and associated *p*-value (*p*_Predictor_). Asterisks indicate predictors that reached statistical significance (non corrected, *p*_Predictor_<0.05). Note that the interaction term was not significant in any network.

| Network | Predictor | $\beta_{\text{Predictor}}$ | S.E. | $t_{\text{Predictor}}(\text{df})$ | $p_{\text{Predictor}}$ |
| --- | --- | --- | --- | --- | --- |
| Parieto-frontal | Manual preference | -0.21 | 0.18 | -1.18 | 0.24 |
|  | Response Hand | -0.24 | 0.17 | -1.36 | 0.17 |
|  | Age (years) | -0.01 | 0.01 | -0.87 | 0.39 |
|  | Education (years) | -0.05 | 0.04 | -1.44 | 0.15 |
|  | T.I.V. (mm<sup>3</sup>) | -1.10<sup>-7</sup> | 7.10<sup>-7</sup> | -0.16 | 0.88 |
|  | Gender | -0.09 | 0.11 | -0.83 | 0.41 |
|  | Language lateralization × Manual preference | 0.16 | 0.17 | 0.94 | 0.35 |
| Posterior-medial | Manual preference | 2.10<sup>-3</sup> | 0.10 | 0.02 | 0.98 |
|  | Response Hand | -0.01 | 0.10 | -0.14 | 0.89 |
|  | Age (years) | -0.01 | 8.10<sup>-3</sup> | -1.41 | 0.16 |
|  | Education (years) | -1.10<sup>-3</sup> | 0.02 | 0.06 | 0.95 |
|  | T.I.V. (mm<sup>3</sup>) | -9.10<sup>-8</sup> | 4.10<sup>-7</sup> | -0.21 | 0.84 |
|  | Gender | -0.09 | 0.06 | -1.42 | 0.16 |
|  | Language lateralization × Manual preference | -0.09 | 0.10 | -0.91 | 0.36 |
| Somato-motor | Manual preference | -0.23 | 0.12 | -1.95 | 0.05 |
|  | Response Hand * | -1.67 | 0.12 | -14.4 | 2.10<sup>-35</sup> |
|  | Age (years) * | -0.03 | 0.01 | -3.19 | 2.10<sup>-3</sup> |
|  | Education (years) | -0.03 | 0.02 | -1.08 | 0.28 |
|  | T.I.V. (mm<sup>3</sup>) | -5.10<sup>-7</sup> | 5.10<sup>-7</sup> | -0.98 | 0.33 |
|  | Gender | 0.11 | 0.07 | 1.48 | 0.14 |
|  | Language lateralization × Manual preference | 0.1 | 0.12 | 0.87 | 0.38 |
| Temporo-frontal | Manual preference | -0.13 | 0.09 | -1.46 | 0.15 |
|  | Response Hand * | -0.21 | 0.09 | -2.46 | 0.01 |
|  | Age (years) | -0.01 | 0.01 | -1.30 | 0.20 |
|  | Education (years) | 1.10<sup>-4</sup> | 0.02 | 6.10<sup>-3</sup> | 1.00 |
|  | T.I.V. (mm<sup>3</sup>) | -7.10<sup>-7</sup> | 4.10<sup>-7</sup> | -1.92 | 0.06 |
|  | Gender | -0.08 | 0.05 | -1.55 | 0.12 |
|  | Language lateralization × Manual preference | 0.13 | 0.08 | 1.56 | 0.12 |
| Visual | Manual preference | -0.09 | 0.11 | -0.79 | 0.43 |
|  | Response Hand | -0.14 | 0.11 | -1.22 | 0.22 |
|  | Age (years) | -0.02 | 0.01 | -2.02 | 0.05 |
|  | Education (years) | 0.03 | 0.02 | 1.08 | 0.28 |
|  | T.I.V. (mm<sup>3</sup>) | 9.10<sup>-7</sup> | 8.10<sup>-7</sup> | 1.92 | 0.06 |
|  | Gender | 0.04 | 0.07 | 0.60 | 0.55 |
|  | Language lateralization × Manual preference | -0.02 | 0.11 | -0.16 | 0.87 |

**Extended Data Table 3.**
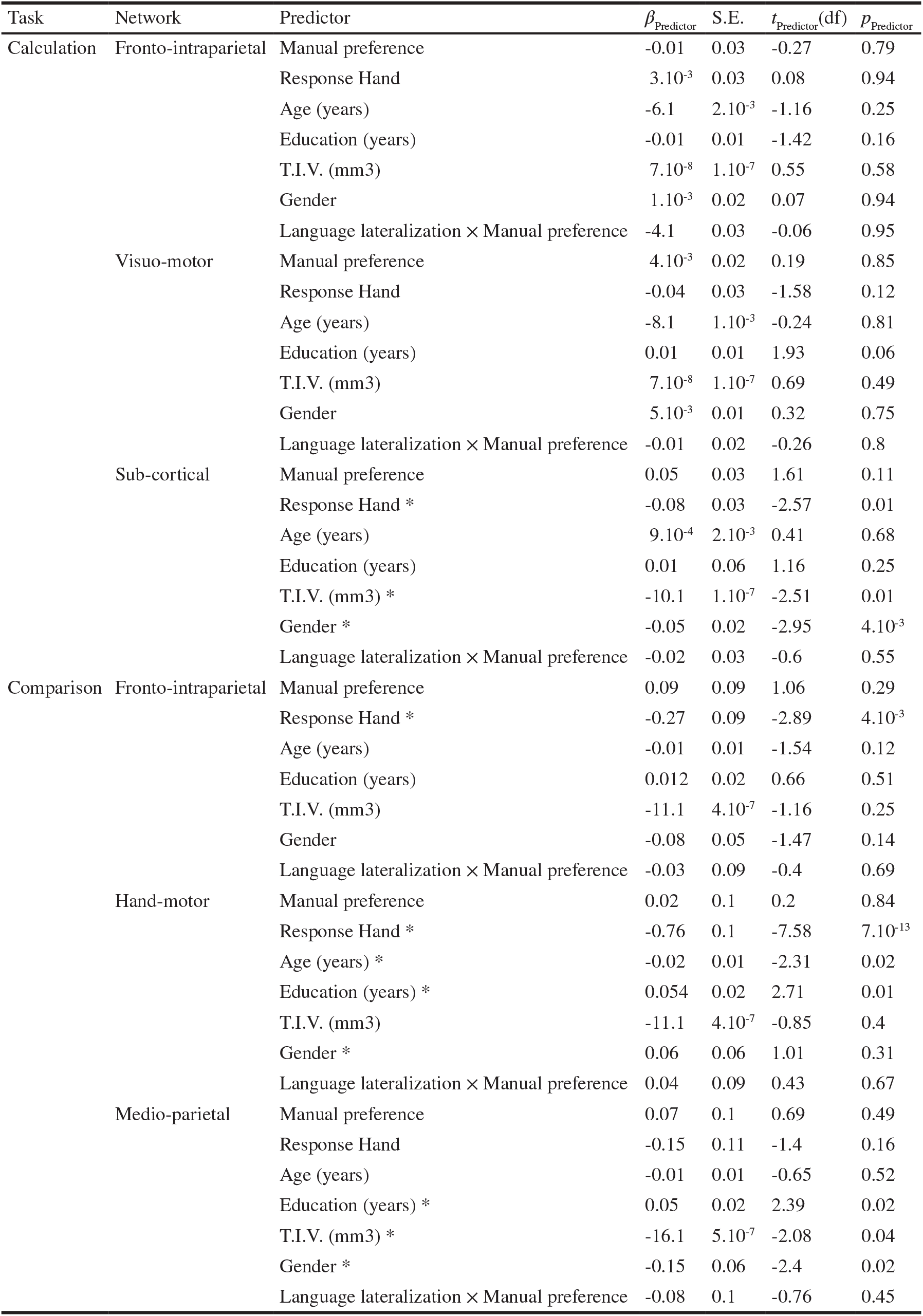
Full regression results for covariates and interaction terms in models assessing the effect of language lateralization phenotype on weighted BOLD asymmetry scores across the six symbolic and analog number networks of LUCA. This table reports the full results for all covariates and the interaction term (language phenotype by manual preference) from the linear regression models presented in Table 3. For each of the six numerical networks, the following are shown for each predictor: regression coefficient (*β*_Predictor_), standard error (S.E.), *t*-value, and associated *p*-value (*p*_Predictor_). Asterisks indicate predictors that reached statistical significance (non corrected, *p*_Predictor_<0.05). Note that the interaction term was not significant in any network.

**Extended Data Table 4.**
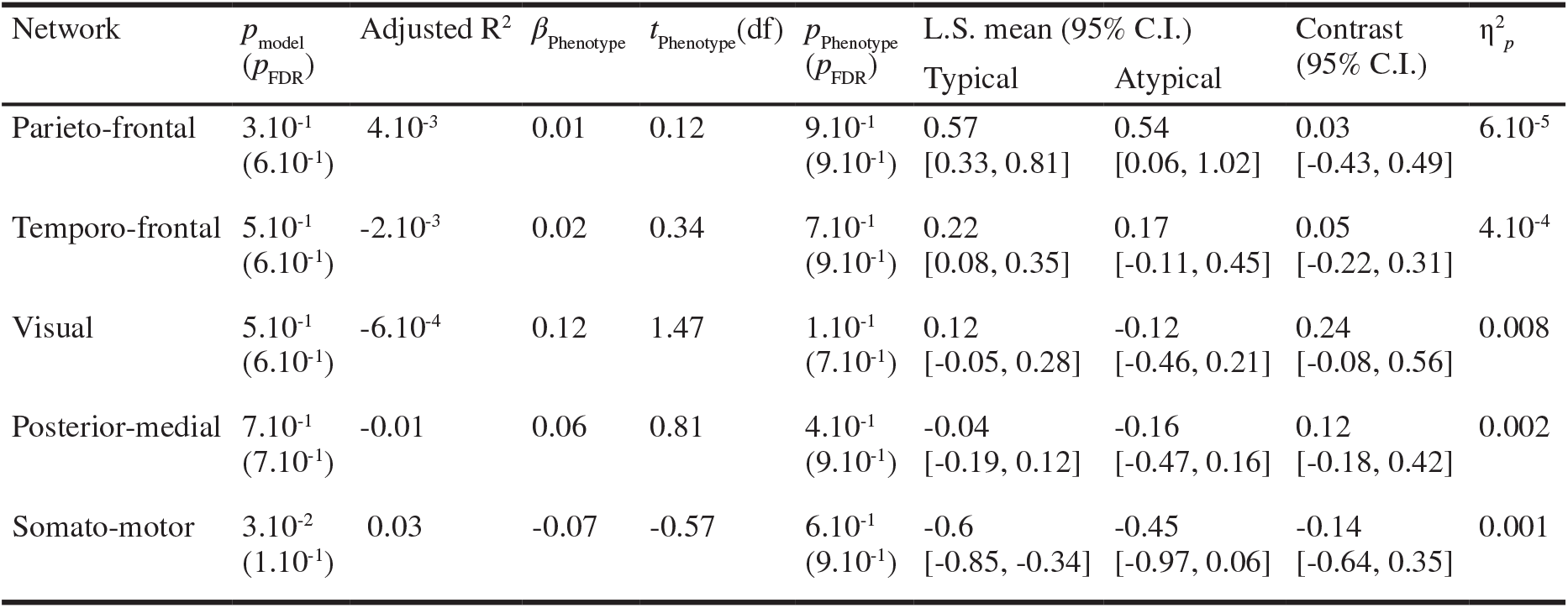
Summary of linear regression results testing the effect of language lateralization phenotype on the degree centrality asymmetry across the five visuospatial attention networks. Separate multiple linear regression models were estimated for each network, with degree centrality asymmetry as the dependent variable. Language lateralization phenotype (typical or atypical) was the primary predictor, controlling for manual preference, age, years of education, total intracranial volume, response hand, and gender. An interaction term between language phenotype and manual preference was also included in all models. The table reports; the overall model p-value (pmodel) and its False Discovery Rate (FDR)-corrected value (*p*_FDR_), the adjusted R^2^, the regression coefficient for language phenotype (*β*_Phenotype_), the *t*-value, and associated *p*-values (uncorrected and FDR-corrected), the least-square means with 95% confidence intervals for typical and atypical groups, the group contrast (difference in estimated means) with 95% confidence interval, and the partial eta-squared (η^2^_*p*_) as a measure of effect size.

**Extended Data Table 5.** Summary of linear regression results testing the effect of language lateralization phenotype on the degree centrality sum across the five visuospatial attention networks. Separate multiple linear regression models were estimated for each network, with degree centrality asymmetry as the dependent variable. Language lateralization phenotype (typical or atypical) was the primary predictor, controlling for manual preference, age, years of education, total intracranial volume, response hand, and gender. An interaction term between language phenotype and manual preference was also included in all models. The table reports; the overall model p-value (pmodel) and its False Discovery Rate (FDR)-corrected value (*p*_FDR_), the adjusted R^2^, the regression coefficient for language phenotype (*β*_Phenotype_), the *t*-value, and associated *p*-values (uncorrected and FDR-corrected), the least-square means with 95% confidence intervals for typical and atypical groups, the group contrast (difference in estimated means) with 95% confidence interval, and the partial eta-squared (η^2^_*p*_) as a measure of effect size.

| Network | $p_{\text{model}}$<br>($p_{\text{FDR}}$) | Adjusted $R^2$ | $\beta_{\text{Phenotype}}$ | $t_{\text{Phenotype}}$ (df) | $p_{\text{Phenotype}}$<br>($p_{\text{FDR}}$) | L.S. mean (95% C.I.) | | Contrast<br>(95% C.I.) | $\eta^2_p$ |
| --- | --- | --- | --- | --- | --- | --- | --- | --- | --- |
|  |  |  |  |  |  | Typical | Atypical |  |  |
| Parieto-frontal | $2 \cdot 10^{-1}$<br>($4 \cdot 10^{-1}$) | 0.01 | 0.15 | 1.53 | $1 \cdot 10^{-1}$<br>($3 \cdot 10^{-1}$) | 4.83<br>[4.63, 5.02] | 4.53<br>[4.14, 4.93] | 0.29<br>[-0.08, 0.67] | 0.008 |
| Temporo-frontal | $2 \cdot 10^{-2}$<br>($1 \cdot 10^{-1}$) | 0.04 | -0.17 | -2.21 | $3 \cdot 10^{-2}$<br>($1 \cdot 10^{-1}$) | 3.53<br>[3.37, 3.69] | 3.87<br>[3.55, 4.19] | -0.34<br>[-0.65, -0.04] | 0.017 |
| Visual | $8 \cdot 10^{-1}$<br>($8 \cdot 10^{-1}$) | -0.01 | 0.04 | 0.31 | $8 \cdot 10^{-1}$<br>($8 \cdot 10^{-1}$) | 4.72<br>[4.43, 5.02] | 4.63<br>[4.04, 5.23] | 0.09<br>[-0.48, 0.66] | $3 \cdot 10^{-4}$ |
| Posterior-medial | $3 \cdot 10^{-1}$<br>($4 \cdot 10^{-1}$) | 0.01 | 0.03 | 0.34 | $7 \cdot 10^{-1}$<br>($8 \cdot 10^{-1}$) | 4.22<br>[4.05, 4.38] | 4.16<br>[3.83, 4.49] | 0.05<br>[-0.26, 0.37] | $4 \cdot 10^{-4}$ |
| Somato-motor | $3 \cdot 10^{-1}$<br>($4 \cdot 10^{-1}$) | $5 \cdot 10^{-3}$ | -0.09 | -0.42 | $7 \cdot 10^{-1}$<br>($8 \cdot 10^{-1}$) | 7.80<br>[7.35, 8.26] | 7.99<br>[7.07, 8.91] | -0.19<br>[-1.07, 0.69] | 0.001 |

**Extended Data Table 6.** Summary of linear regression results testing the effect of language lateralization phenotype on the inter-hemispheric homotopic intrinsic correlation (Fisher *z*-transformed) across the five visuospatial attention networks. Separate multiple linear regression models were estimated for each network, with degree centrality asymmetry as the dependent variable. Language lateralization phenotype (typical or atypical) was the primary predictor, controlling for manual preference, age, years of education, total intracranial volume, response hand, and gender. An interaction term between language phenotype and manual preference was also included in all models. The table reports; the overall model p-value (pmodel) and its False Discovery Rate (FDR)-corrected value (*p*_FDR_), the adjusted R^2^, the regression coefficient for language phenotype (*β*_Phenotype_), the *t*-value, and associated *p*-values (uncorrected and FDR-corrected), the least-square means with 95% confidence intervals for typical and atypical groups, the group contrast (difference in estimated means) with 95% confidence interval, and the partial eta-squared (η^2^_*p*_) as a measure of effect size.

| Network | $p_{\text{model}}$<br>($p_{\text{FDR}}$) | Adjusted $R^2$ | $\beta_{\text{Phenotype}}$ | $t_{\text{Phenotype}}$ (df) | $p_{\text{Phenotype}}$<br>($p_{\text{FDR}}$) | L.S. mean (95% C.I.) | | Contrast<br>(95% C.I.) | $\eta_p^2$ |
| --- | --- | --- | --- | --- | --- | --- | --- | --- | --- |
|  |  |  |  |  |  | Typical | Atypical |  |  |
| Parieto-frontal | $6.10^{-1}$<br>($6.10^{-1}$) | $-4.10^{-3}$ | -0.02 | -0.57 | $6.10^{-1}$<br>($6.10^{-1}$) | 1.29<br>[1.22, 1.37] | 1.33<br>[1.18, 1.49] | -0.04<br>[-0.19, 0.10] | $1.10^{-3}$ |
| Temporo-frontal | $3.10^{-1}$<br>($3.10^{-1}$) | 0.01 | -0.02 | -0.85 | $4.10^{-1}$<br>($5.10^{-1}$) | 0.83<br>[0.77, 0.88] | 0.87<br>[0.76, 0.99] | -0.05<br>[-0.16, 0.06] | 0.003 |
| Visual | $2.10^{-1}$<br>($3.10^{-1}$) | 0.01 | -0.04 | -1.18 | $2.10^{-1}$<br>($4.10^{-1}$) | 1.39<br>[1.33, 1.45] | 1.46<br>[1.33, 1.59] | -0.07<br>[-0.20, 0.05] | 0.005 |
| Posterior-medial | $2.10^{-3}$<br>($8.10^{-3}$) | 0.06 | -0.05 | -1.98 | $5.10^{-2}$<br>($2.10^{-1}$) | 1.09<br>[1.04, 1.15] | 1.20<br>[1.09, 1.30] | -0.10<br>[-0.20, -0.00] | 0.014 |
| Somato-motor | $2.10^{-1}$<br>($3.10^{-1}$) | 0.01 | -0.05 | -1.78 | $8.10^{-2}$<br>($2.10^{-1}$) | 0.95<br>[0.89, 1.01] | 1.05<br>[0.93, 1.17] | -0.10<br>[-0.21, 0.01] | 0.011 |

**Extended Data Table 7.** Summary of linear regression results testing the effect of language lateralization phenotype on the degree of centrality asymmetry across the six lateralized networks supporting arithmetics and numerical comparisons of LUCA. Separate multiple linear regression models were estimated for each network, with degree centrality asymmetry as the dependent variable. Language lateralization phenotype (typical or atypical) was the primary predictor, controlling for manual preference, age, years of education, total intracranial volume, response hand, and gender. An interaction term between language phenotype and manual preference was also included in all models. The table reports; the overall model *p*-value (*p*_model_) and its False Discovery Rate (FDR)-corrected value (*p*_FDR_), the adjusted R^2^, the regression coefficient for language phenotype (*β*_Phenotype_), the *t*-value, and associated *p*-values (uncorrected and FDR-corrected), the least-square means with 95% confidence intervals for typical and atypical groups, the group contrast (difference in estimated means) with 95% confidence interval, and the partial eta-squared (η^2^_*p*_) as a measure of effect size.

| Task | Network | $p_{\text{model}}$<br>($p_{\text{FDR}}$) | Adjusted $R^2$ | $\beta_{\text{Phenotype}}$ | $t_{\text{Phenotype}}$ (df) | $p_{\text{Phenotype}}$<br>($p_{\text{FDR}}$) | L.S. mean (95% C.I.) | | Contrast<br>(95% C.I.) | $\eta_p^2$ |
| --- | --- | --- | --- | --- | --- | --- | --- | --- | --- | --- |
|  |  |  |  |  |  |  | Typical | Atypical |  |  |
| Calculation | Fronto-intraparietal | $8 \cdot 10^{-1}$<br>($9 \cdot 10^{-1}$) | -0.012 | -0.02 | -0.33 | $7 \cdot 10^{-1}$<br>($9 \cdot 10^{-1}$) | 0.00<br>[-0.06, 0.06] | 0.04<br>[-0.20, 0.28] | -0.04<br>[-0.29, 0.20] | $4 \cdot 10^{-4}$ |
| | Visuo-motor | $3 \cdot 10^{-1}$<br>($6 \cdot 10^{-1}$) | 0.01 | 0.05 | 0.80 | $4 \cdot 10^{-1}$<br>($9 \cdot 10^{-1}$) | -0.03<br>[-0.09, 0.03] | -0.14<br>[-0.38, 0.11] | 0.10<br>[-0.15, 0.36] | 0.002 |
| | Sub-cortical | $6 \cdot 10^{-1}$<br>($9 \cdot 10^{-1}$) | -0.01 | 0.03 | 0.38 | $7 \cdot 10^{-1}$<br>($9 \cdot 10^{-1}$) | 0.40<br>[0.33, 0.46] | 0.35<br>[0.09, 0.60] | 0.05<br>[-0.21, 0.32] | 0.001 |
| Comparison | Fronto-intraparietal | $6 \cdot 10^{-2}$<br>($3 \cdot 10^{-1}$) | 0.02 | 0.01 | 0.13 | $9 \cdot 10^{-1}$<br>($9 \cdot 10^{-1}$) | 0.41<br>[0.35, 0.48] | 0.40<br>[0.14, 0.65] | 0.02<br>[-0.24, 0.28] | $6 \cdot 10^{-5}$ |
| | Hand-motor | $2 \cdot 10^{-1}$<br>($6 \cdot 10^{-1}$) | 0.01 | 0.09 | 1.41 | $2 \cdot 10^{-1}$<br>($9 \cdot 10^{-1}$) | -0.03<br>[-0.09, 0.04] | -0.21<br>[-0.45, 0.04] | 0.18<br>[-0.07, 0.43] | 0.007 |
| | Medio-parietal | $9 \cdot 10^{-1}$<br>($9 \cdot 10^{-1}$) | -0.02 | -0.02 | -0.42 | $7 \cdot 10^{-1}$<br>($9 \cdot 10^{-1}$) | -0.05<br>[-0.10, 0.00] | -0.01<br>[-0.20, 0.18] | -0.04<br>[-0.24, 0.15] | 0.001 |

**Extended Data Table 8.**
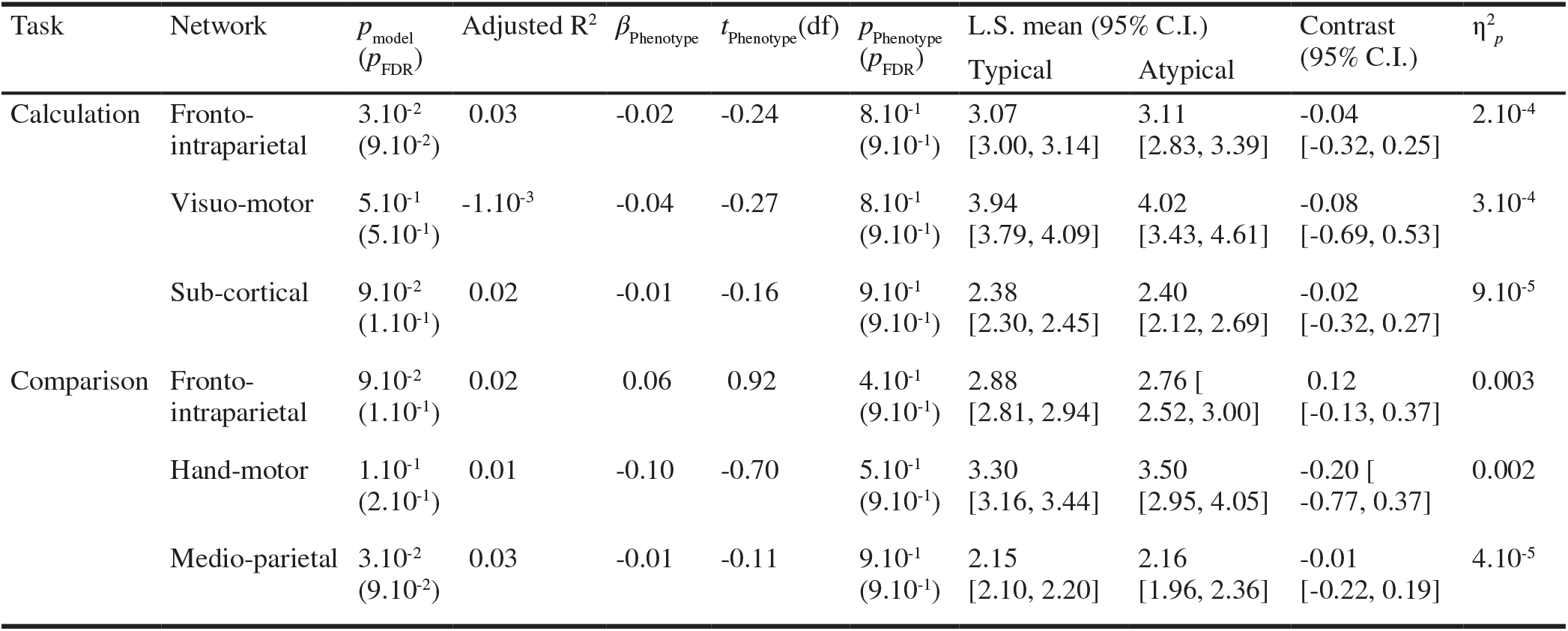
Summary of linear regression results testing the effect of language lateralization phenotype on the degree of centrality sum across the six lateralized networks supporting arithmetics and numerical comparisons of LUCA. Separate multiple linear regression models were estimated for each network, with degree centrality asymmetry as the dependent variable. Language lateralization phenotype (typical or atypical) was the primary predictor, controlling for manual preference, age, years of education, total intracranial volume, response hand, and gender. An interaction term between language phenotype and manual preference was also included in all models. The table reports; the overall model *p*-value (*p*_model_) and its False Discovery Rate (FDR)-corrected value (*p*_FDR_), the adjusted R^2^, the regression coefficient for language phenotype (*β*_Phenotype_), the *t*-value, and associated *p*-values (uncorrected and FDR-corrected), the least-square means with 95% confidence intervals for typical and atypical groups, the group contrast (difference in estimated means) with 95% confidence interval, and the partial eta-squared (η^2^_*p*_) as a measure of effect size.

**Extended Data Table 9.** Summary of linear regression results testing the effect of language lateralization phenotype on the interhemispheric homotopic intrinsic correlation (Fisher *z*-transformed) across the six lateralized networks supporting arithmetics and numerical comparisons of LUCA. Separate multiple linear regression models were estimated for each network, with degree centrality asymmetry as the dependent variable. Language lateralization phenotype (typical or atypical) was the primary predictor, controlling for manual preference, age, years of education, total intracranial volume, response hand, and gender. An interaction term between language phenotype and manual preference was also included in all models. The table reports; the overall model *p*-value (*p*_model_) and its False Discovery Rate (FDR)-corrected value (*p*_FDR_), the adjusted R^2^, the regression coefficient for language phenotype (*β*_Phenotype_), the *t*-value, and associated *p*-values (uncorrected and FDR-corrected), the least-square means with 95% confidence intervals for typical and atypical groups, the group contrast (difference in estimated means) with 95% confidence interval, and the partial eta-squared (η^2^_*p*_) as a measure of effect size.

| Task | Network | $p_{\text{model}}$<br>($p_{\text{FDR}}$) | Adjusted $R^2$ | $\beta_{\text{Phenotype}}$ | $t_{\text{Phenotype}}$(df) | $p_{\text{Phenotype}}$<br>($p_{\text{FDR}}$) | L.S. mean (95% C.I.) | | Contrast<br>(95% C.I.) | $\eta_p^2$ |
| --- | --- | --- | --- | --- | --- | --- | --- | --- | --- | --- |
|  |  |  |  |  |  |  | Typical | Atypical |  |  |
| Calculation | Fronto-intraparietal | $6.10^{-3}$<br>($3.10^{-2}$) | 0.04 | -0.01 | -1.81 | $7.10^{-2}$<br>($1.10^{-1}$) | 0.28<br>[0.27, 0.28] | 0.30<br>[0.27, 0.33] | -0.02<br>[-0.05, 0.00] | 0.012 |
| | Visuo-motor | $8.10^{-2}$<br>($1.10^{-1}$) | 0.02 | -0.02 | -1.86 | $6.10^{-2}$<br>($1.10^{-1}$) | 0.64<br>[0.63, 0.65] | 0.67<br>[0.64, 0.71] | -0.04<br>[-0.07, 0.00] | 0.012 |
| | Sub-cortical | $1.10^{-1}$<br>($1.10^{-1}$) | 0.02 | -0.01 | -0.99 | $3.10^{-1}$<br>($3.10^{-1}$) | 0.69<br>[0.68, 0.70] | 0.71<br>[0.67, 0.75] | -0.02<br>[-0.06, 0.02] | 0.004 |
| Comparison | Fronto-intraparietal | $1.10^{-1}$<br>($1.10^{-1}$) | 0.02 | -0.01 | -1.16 | $2.10^{-1}$<br>($3.10^{-1}$) | 0.19<br>[0.18, 0.20] | 0.20<br>[0.18, 0.23] | -0.01<br>[-0.04, 0.01] | 0.005 |
| | Hand-motor | $1.10^{-1}$<br>($1.10^{-1}$) | 0.02 | -0.01 | -1.27 | $2.10^{-1}$<br>($3.10^{-1}$) | 0.66<br>[0.65, 0.67] | 0.69<br>[0.65, 0.73] | -0.03<br>[-0.07, 0.02] | 0.006 |
| | Medio-parietal | $9.10^{-3}$<br>($3.10^{-2}$) | 0.04 | -0.01 | -1.79 | $7.10^{-2}$<br>($1.10^{-1}$) | 0.74<br>[0.74, 0.75] | 0.77<br>[0.74, 0.79] | -0.02<br>[-0.05, 0.00] | 0.011 |

## References

1. A. W. Toga, P. M. Thompson, Mapping brain asymmetry. Nat Rev Neurosci 4, 37–48 (2003).

2. M. C. Corballis, The evolution of language. Ann N Y Acad Sci 1156, 19–43 (2009).

3. P.-Y. Hervé, L. Zago, L. Petit, B. Mazoyer, N. Tzourio-Mazoyer, Revisiting human hemispheric specialization with neuroimaging. Trends Cogn Sci 17, 69–80 (2013).

4. G. Ojemann, J. Ojemann, E. Lettich, M. Berger, Cortical language localization in left, dominant hemisphere. An electrical stimulation mapping investigation in 117 patients. J Neurosurg 71, 316–326 (1989).

5. J. Wada, T. Rasmussen, Intracarotid injection of sodium amytal for the lateralization of cerebral speech dominance. J. eurosurg. 17, 266–282 (1960).

6. N. Dronkers, J. Ogar, Brain areas involved in speech production. Brain 127, 1461–1462 (2004).

7. C. J. Price, The anatomy of language: contributions from functional neuroimaging. J Anat 197 Pt 3, 335–359 (2000).

8. G. Vingerhoets, Phenotypes in hemispheric functional segregation? Perspectives and challenges. Phys Life Rev 30, 1–18 (2019).

9. M. Corbetta, G. L. Shulman, Control of goal-directed and stimulus-driven attention in the brain. Nat Rev Neurosci 3, 201–215 (2002).

10. H.-O. Karnath, C. Rorden, The anatomy of spatial neglect. Neuropsychologia 50, 1010–1017 (2012).

11. M. Kinsbourne, The cerebral basis of lateral asymmetries in attention. Acta Psychol (Amst) 33, 193–201 (1970).

12. M. M. Mesulam, Spatial attention and neglect: parietal, frontal and cingulate contributions to the mental representation and attentional targeting of salient extrapersonal events. Philos Trans R Soc Lond B Biol Sci 354, 1325–1346 (1999).

13. N. Kanwisher, J. McDermott, M. M. Chun, The fusiform face area: a module in human extrastriate cortex specialized for face perception. J Neurosci 17, 4302–4311 (1997).

14. Q. Cai, L. Van der Haegen, M. Brysbaert, Complementary hemispheric specialization for language production and visuospatial attention. Proc Natl Acad Sci U S A 110, E322–30 (2013).

15. N. Tzourio-Mazoyer, L. Zago, B. Mazoyer, What can we learn from healthy atypical individuals on the segregation of complementary functions?: Comment on “Phenotypes in hemispheric functional segregation? Perspectives and challenges” by Guy Vingerhoets. Phys Life Rev 30, 34–37 (2019).

16. L. Labache, et al., A SENtence Supramodal Areas AtlaS (SENSAAS) based on multiple task-induced activation mapping and graph analysis of intrinsic connectivity in 144 healthy right-handers. Brain Struct. Funct. 224, 859–882 (2019).

17. I. Hesling, L. Labache, M. Joliot, N. Tzourio-Mazoyer, Large-scale plurimodal networks common to listening to, producing and reading word lists: an fMRI study combining task-induced activation and intrinsic connectivity in 144 right-handers. Brain Struct. Funct. 224, 3075–3094 (2019).

18. L. Labache, L. Petit, M. Joliot, L. Zago, Atlas for the Lateralized Visuospatial Attention Networks (ALANs): Insights from fMRI and network analyses. Imaging Neuroscience 2, 1–22 (2024).

19. P. Pinel, M. Piazza, D. Le Bihan, S. Dehaene, Distributed and overlapping cerebral representations of number, size, and luminance during comparative judgments. Neuron 41, 983–993 (2004).

20. L. Zago, et al., How verbal and spatial manipulation networks contribute to calculation: an fMRI study. Neuropsychologia 46, 2403–2414 (2008).

21. P. Pinel, S. Dehaene, Beyond hemispheric dominance: brain regions underlying the joint lateralization of language and arithmetic to the left hemisphere. J Cogn Neurosci 22, 48–66 (2010).

22. L. Zago, et al., Neural correlates of simple and complex mental calculation. Neuroimage 13, 314–327 (2001).

23. N. Tzourio-Mazoyer, L. Zago, H. Cochet, F. Crivello, Development of handedness, anatomical and functional brain lateralization. Handb Clin Neurol 173, 99–105 (2020).

24. N. Tzourio-Mazoyer, M. Perrone-Bertolotti, G. Jobard, B. Mazoyer, M. Baciu, Multi-factorial modulation of hemispheric specialization and plasticity for language in healthy and pathological conditions: A review. Cortex 86, 314–339 (2017).

25. P. Pica, C. Lemer, V. Izard, S. Dehaene, Exact and approximate arithmetic in an Amazonian indigene group. Science 306, 499–503 (2004).

26. R. Gerrits, Variability in Hemispheric Functional Segregation Phenotypes: A Review and General Mechanistic Model. Neuropsychol Rev 34, 27–40 (2024).

27. D. J. Serrien, L. O’Regan, The interactive functional biases of manual, language and attention systems. Cogn Res Princ Implic 7, 20 (2022).

28. M. S. Gazzaniga, Cerebral specialization and interhemispheric communication: does the corpus callosum enable the human condition? Brain 123 (Pt 7), 1293–1326 (2000).

29. L. Labache, S. Chopra, X.-H. Zhang, A. J. Holmes, The molecular and cellular underpinnings of human brain lateralization. bioRxiv (2025).

30. R. Rajimehr, A. Firoozi, H. Rafipoor, N. Abbasi, J. Duncan, Complementary hemispheric lateralization of language and social processing in the human brain. Cell Rep 41, 111617 (2022).

31. N. Tzourio-Mazoyer, L. Labache, L. Zago, I. Hesling, B. Mazoyer, Neural support of manual preference revealed by BOLD variations during right and left finger-tapping in a sample of 287 healthy adults balanced for handedness. Laterality 1–23 (2021).

32. S. Ocklenburg, et al., Clinical implications of brain asymmetries. Nat Rev Neurol 20, 383–394 (2024).

33. R. Gerrits, H. Verhelst, G. Vingerhoets, Mirrored brain organization: Statistical anomaly or reversal of hemispheric functional segregation bias? Proc Natl Acad Sci U S A 117, 14057–14065 (2020).

34. E. Villar-Rodríguez, T. Davydova, L. Marin-Marin, C. Avila, Atypical lateralization of visuospatial attention can be associated with better or worse performance on line bisection. Brain Struct Funct 229, 1577–1590 (2024).

35. L. Labache, et al., Typical and atypical language brain organization based on intrinsic connectivity and multitask functional asymmetries. Elife 9 (2020).

36. L. Labache, T. Ge, B. Yeo, A. Holmes, Language network lateralization is reflected throughout the macroscale functional organization of cortex. Nat. Commun. 14, 3405 (2023).

37. B. Mazoyer, et al., BIL&GIN: A neuroimaging, cognitive, behavioral, and genetic database for the study of human brain lateralization. Neuroimage 124, 1225–1231 (2016).

38. B. Mazoyer, et al., BIL&GIN: A neuroimaging, cognitive, behavioral, and genetic database for the study of human brain lateralization. Neuroimage 124, 1225–1231 (2016).

39. L. Zago, et al., The association between hemispheric specialization for language production and for spatial attention depends on left-hand preference strength. Neuropsychologia 93, 394–406 (2016).

40. G. Josse, N. Tzourio-Mazoyer, Hemispheric specialization for language. Brain Res Brain Res Rev 44, 1–12 (2004).

41. N. Tzourio-Mazoyer, C. Courtin, “Brain lateralization and the emergence of language” in Studies in Language Companion Series, (John Benjamins Publishing Company, 2013), pp. 237–256.

42. N. Tzourio-Mazoyer, M. L. Seghier, The neural bases of hemispheric specialization. Neuropsychologia 93, 319–324 (2016).

43. J. E. Quin-Conroy, D. M. Bayliss, S. G. Daniell, N. A. Badcock, Patterns of language and visuospatial functional lateralization and cognitive ability: a systematic review. Laterality 29, 63–96 (2024).

44. R. Rolinski, et al., Language lateralization from task-based and resting state functional MRI in patients with epilepsy. Hum. Brain Mapp. 41, 3133–3146 (2020).

45. A. Teghipco, A. Hussain, M. E. Tivarus, Disrupted functional connectivity affects resting state based language lateralization. Neuroimage Clin 12, 910–927 (2016).

46. C. Semenza, S. Benavides-Varela, E. Salillas, Brain laterality of numbers and calculation: Complex networks and their development. Handb Clin Neurol 208, 461–480 (2025).

47. C. Cano-Melle, E. Villar-Rodríguez, M. Baena-Pérez, M. arcet, C. Avila, Effects of Lateralization of Language on Cognition Among Left-Handers. Neurobiol Lang (Camb) 6 (2025).

48. T. R. J. Gonzalez Alam, et al., A double dissociation between semantic and spatial cognition in visual to default network pathways. Elife 13 (2025).

49. S. Kress, J. Neudorf, C. Ekstrand, R. Borowsky, Exploring the interaction of reading and attention through connectivity with the frontal-eye-field. Neuroscience 585, 249–261 (2025).

50. S. J. Gotts, et al., Two distinct forms of functional lateralization in the human brain. Proc Natl Acad Sci U S A 110, E3435–44 (2013).

51. B. T. T. Yeo, et al., Functional Specialization and Flexibility in Human Association Cortex. Cereb Cortex 25, 3654–3672 (2015).

52. M. Joliot, et al., AICHA: An atlas of intrinsic connectivity of homotopic areas. J. Neurosci. Methods 254, 46–59 (2015).

53. R Core Team, R: A language and environment for statistical computing (R Foundation for Statistical Computing, 2021).

54. J. Fox, S. Weisberg, B. Price, Car: Companion to applied regression. The R Foundation. 10.32614/cran.package.car. Deposited 1 May 2001.

55. H. Wickham, R. François, L. Henry, K. Müller, D. Vaughan, dplyr: A Grammar of Data Manipulation. The R Foundation. 10.32614/cran.package.dplyr. Deposited 16 January 2014.

56. H. Wickham, D. Vaughan, M. Girlich, tidyr: Tidy Messy Data. The R Foundation. 10.32614/cran.package.tidyr. Deposited 21 July 2014.

57. H. Wickham, L. Henry, purrr: Functional Programming Tools. The R Foundation. 10.32614/cran.package.purrr. Deposited 28 September 2015.

58. D. Robinson, A. Hayes, S. Couch, broom: Convert Statistical Objects into Tidy Tibbles. The R Foundation. 10.32614/cran.package.broom. Deposited 23 November 2014.

59. R. V. Lenth, emmeans: Estimated Marginal Means, aka Least-Squares Means. The R Foundation. 10.32614/cran.package.emmeans. Deposited 20 October 2017.

60. M. S. Ben-Shachar, et al., Effectsize: Indices of effect size. The R Foundation. 10.32614/cran.package.effectsize. Deposited 15 November 2019.

61. H. Wickham, et al., Ggplot2: Create elegant data visualisations using the grammar of graphics. The R Foundation. 10.32614/cran.package.ggplot2. Deposited 1 June 2007.

62. NITRC: Surf ice: Tool/resource info. Available at: http://www.nitrc.org/projects/surfice/ [Accessed 24 March 2022].

63. C. Rorden, Surfice: visualizing neuroimaging meshes, tractography streamlines and connectomes. Nat Methods (2025). 10.1038/s41592-025-02764-6.

64. S. Chopra, L. Labache, E. Dhamala, E. R. Orchard, A. Holmes, A practical guide for generating reproducible and programmatic neuroimaging visualizations. Aperture Neuro 3 (2023).

65. L. Labache, et al., Typical and atypical language brain organization based on intrinsic connectivity and multitask functional asymmetries. Elife 9, e58722 (2020).

66. Y. Benjamini, Y. Hochberg, Controlling the false discovery rate: A practical and powerful approach to multiple testing. J. R. Stat. Soc. Series B Stat. Methodol. 57, 289–300 (1995).

67. L. Labache, loiclabache/ALANs_brainAtlas: Atlas for the Lateralized Visuospatial Attention Networks (ALANs) (Zenodo, 2024).

